# Heterogeneous NF-κB activation and enhancer features shape transcription in *Drosophila* immunity

**DOI:** 10.1101/2025.05.19.654881

**Authors:** Noshin Nawar, Emma Rits, Lianne Cohen, Zeba Wunderlich

## Abstract

Conserved NF-κB signaling pathways shape immune responses in animals. In mammals, NF-κB activation patterns and downstream transcription vary with stimulus, cell type, and stochastic differences among identically treated cells. Whether animals without adaptive immunity exhibit similar heterogeneity or rely on distinct immune strategies remains unknown. We engineered *Drosophila melanogaster* S2* reporter cells as an immune-responsive model to monitor the dynamics of an NF-κB transcription factor, Relish, and downstream transcription in single, living cells. Following immune stimulation, Relish exhibits diverse nuclear localization dynamics that fall into distinct categories, with both the fraction of responsive cells and their activation speed rising with stimulus dose. Pre-stimulus features, including Relish nuclear fraction, predict a cell’s responsiveness to stimulation. Simultaneous measurement of Relish and downstream transcription revealed that the probability of transcriptional bursts from immune-responsive enhancers correlates with Relish nuclear fraction. The number of NF-κB binding sites tunes transcriptional activity among immune enhancers. Our study uncovers heterogeneity in NF-κB activation and target gene expression within *Drosophila*, illustrating how dynamic NF-κB behavior and enhancer architecture tune gene regulation.

## Introduction

Across the animal kingdom, both vertebrate and invertebrate metazoans possess an innate immune system to combat pathogenic threats. This evolutionarily conserved defense mechanism relies on germline-encoded receptors that recognize pathogen-associated molecular patterns and trigger the nuclear translocation of NF-κB transcription factors. Nuclear NF-κB binds κB motifs in enhancers to induce the expression of immune effector genes, e.g., antimicrobial peptides (AMPs) ^1–4^. These pathways facilitate rapid pathogen recognition and elimination, conferring resistance to infections.

In mammals, extensive work has demonstrated that NF-κB activation is not a uniform “on/off” switch, but rather is regulated by diverse signaling dynamics that vary depending on cell type and stimulus. For instance, fibroblasts exhibit oscillatory nuclear-cytoplasmic shuttling of NF-κB, while macrophages display sustained or transient activation ^5–8^. Among genetically identical populations of cells, stochastic fluctuations have been observed in which some cells rapidly initiate robust signaling, and others show little to no response ^5–9^. Heterogeneity in signaling is essential, allowing mammalian cells to respond efficiently and appropriately to diverse stimuli while avoiding excessive inflammation ^10,11^.

While NF-κB transcription factors are found in nearly all eukaryotic genomes from protists to mammals — though notably absent in nematodes and yeast — the dynamics and consequences of their activation remain poorly understood outside of vertebrates ^12^. Most animals lack an adaptive immune system, and it remains unclear whether these organisms exhibit heterogeneity similar to mammals or rely on distinct immune strategies. The fruit fly *Drosophila melanogaster* is a powerful model for dissecting innate immunity due to its evolutionary conservation of key signaling pathways and transcriptional regulators. Its lack of adaptive immunity allows for specific investigation into innate immunity without confounding adaptive responses, and flies can be studied as an integrated system at the level of the whole organism. *Drosophila*’s hallmark innate immune response to microbes is regulated by two distinct NF-κB signaling mechanisms: the immune deficiency (IMD) and Toll pathways (Figure 1A). IMD is highly specialized to detect pathogen-derived meso-diaminopimelic acid (DAP)-type peptidoglycans commonly found on the surface of Gram-negative bacteria and certain Gram-positive bacteria, such as *Bacillus* and *Listeria* ^3,13,14^. Upon recognition, the NF-κB Relish translocates to the nucleus to facilitate rapid AMP induction in the fat body and hemocytes ^13,15^. Robust AMP gene expression has served as a convenient readout for pathway activity.

**Figure 1.**
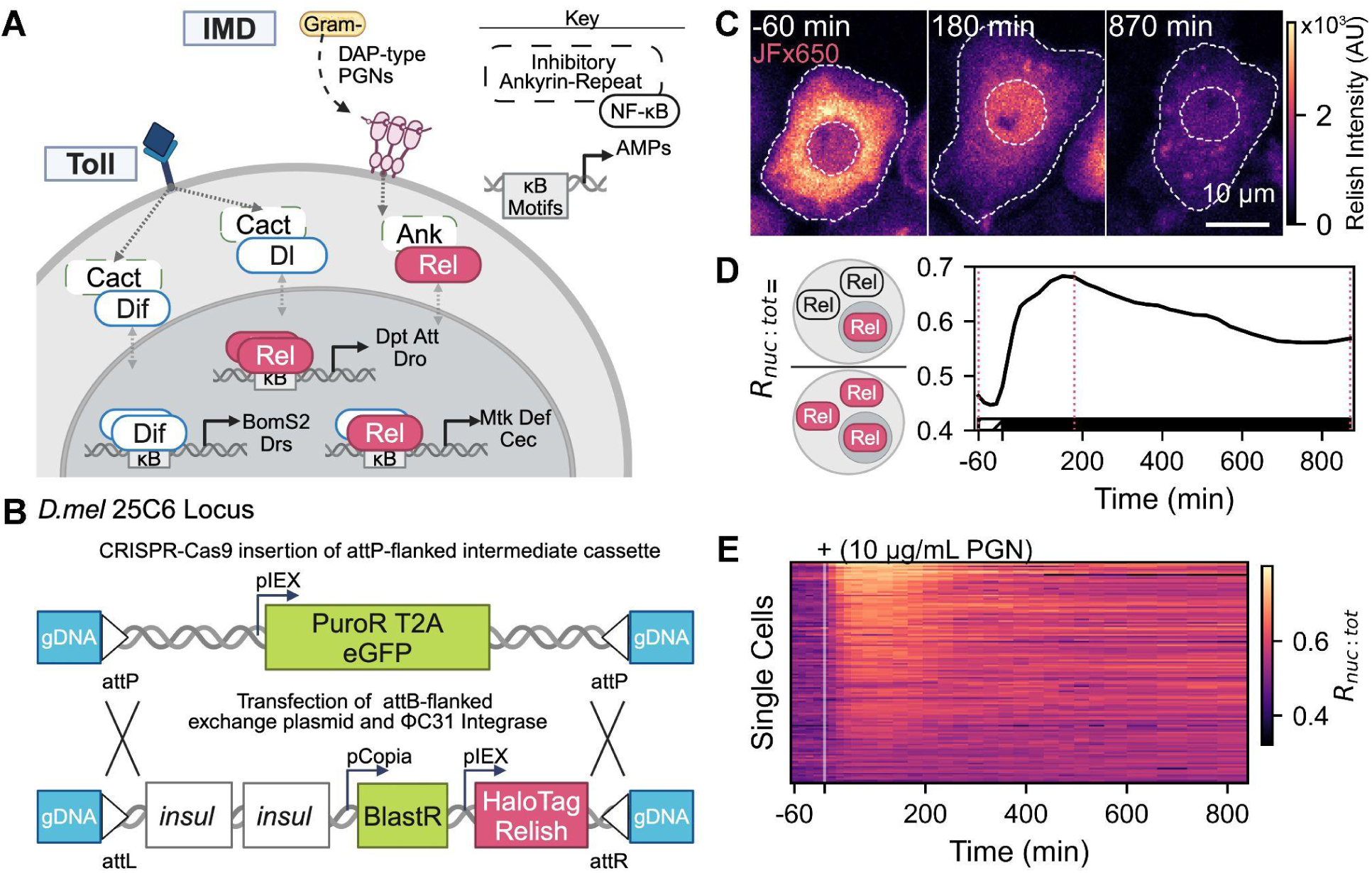
Spatiotemporal monitoring of IMD pathway activation using an S2* reporter cell line. **A.** Schematic of the *Drosophila* humoral innate immune response. Gram-positive bacteria, fungi and yeast activate the Toll pathway via proteolytic cleavage and nuclear translocation of NF-κB transcription factors Dif and Dorsal (Dl) ^3,14,31^. Gram-negative bacteria are detected by peptidoglycan-recognition proteins in the IMD pathway, triggering Relish cleavage and nuclear translocation into the nucleus where NF-κB homodimers and heterodimers bind κB enhancer motifs to drive antimicrobial peptide (AMP) gene expression. **B.** Schematic illustrating CRISPR/Cas9-mediated insertion of an attP-flanked repair cassette at the 25C6 locus, followed by ΦC31 recombinase-mediated cassette exchange (RMCE) to introduce gypsy insulators (insul), blasticidin resistance (BlastR), and HaloTag–Relish flanked by attL/R sites ^29^. **C.** A representative cell’s JFx650-bound HaloTag-Relish signal before (left) and after (middle, right) addition of 10 µg/mL PGN. Overlaid masks show cell bodies and nuclei. Images are confocal maximum-intensity projections (three z-steps spanning 2 µm). Scale bar = 10 µm. **D.** (Left) Schematic representing R_nuc:tot_, the ratio of nuclear to total HaloTag-Relish signal, and its time course (right) for the cell in C. Horizontal bar on the x-axis denotes the pre- (hatched) and post- (solid) injection phases. Vertical lines mark selected timepoints in C. Data were smoothed using Savitzky Golay filtering with a polynomial order of 2. **E.** Heatmap of R_nuc:tot_ over time in response to PGN injection at time = 0 (n = 129 cells (rows) from 4 independent replicates). Schematics in A and B were created using BioRender (https://biorender.com).

Most studies of AMP induction in *Drosophila* have relied on bulk measurements of transcription across tissues or whole flies, obscuring potential cell-to-cell variability that may be critical for understanding regulation ^16–18^. Recent single-cell investigations of the fat body and hemocytes have begun to reveal that AMP expression is restricted to a subset of cells during immune challenges such as sterile wounding, bacterial infection, or parasitic wasp infection ^19–22^. Among responsive cells, AMP expression levels may vary. These studies, though informative, are limited to fixed time points, providing limited insight into the temporal dynamics of AMP expression and, crucially, the upstream regulation by NF-κB driving transcriptional variability. In particular, little is known about how the spatiotemporal behavior of the NF-κB Relish contributes to differential AMP expression at the single-cell level.

To address these gaps, we engineered live reporters in immune-responsive *Drosophila* S2* cells to concurrently track Relish dynamics and AMP transcription. This approach enabled us to quantify the relationship between upstream Relish spatiotemporal behavior and downstream transcriptional responses at single-cell resolution, identifying where variability arises in the IMD signaling pathway. Using a HaloTag-labeled Relish ^23^ reporter line, we identified distinct categories of Relish nuclear translocation dynamics following stimulation, with higher stimulus levels increasing both the fraction of responsive cells and the immediacy of the response. A cell’s likelihood of activation was linked to pre-stimulus features, including its baseline nuclear Relish level. Leveraging recent advances in RNA aptamer imaging ^24^, we built upon our Relish reporter to enable live monitoring of transcription driven by enhancers of key antimicrobial peptides alongside upstream Relish signaling. Higher levels of nuclear Relish were a strong predictor of transcriptional activity, and the proportion of transcriptionally responsive cells was tightly, though non-linearly, coupled to nuclear Relish levels. The ratio of transcriptionally active cells in the population was modulated by AMP enhancer architecture. High-frequency imaging further revealed enhancer-specific control of transcriptional burst kinetics. Together, these findings uncover novel layers of heterogeneity in immune signaling and gene expression, offering key insights into how signaling dynamics drive functional heterogeneity in immune responses, ultimately shaping the effectiveness of organismal immunity ^9,21,22,25,26^.

## Results

### Immune-responsive S2* cells display heterogeneous NF-κB dynamics upon IMD stimulation

To investigate NF-κB dynamics across an isogenic cell population, we employed hemocyte-like, late-stage embryo-derived *Drosophila* S2* cells for their robust IMD response ^27,28^. NF-κB subcellular localization was visualized with time-lapse confocal imaging of Relish fused with an N-terminal HaloTag ^23^. We inserted a single copy of the transgene cassette at the 25C6 locus of S2* cells via recombination mediated cassette exchange (RMCE), reducing expression variability across the cell population ^29^ (Figure 1B). Polymerase chain reaction and sequencing of the resulting cells’ loci confirmed the insertion of the full transgene (Figure S1).

Relish spatiotemporal dynamics were imaged across 1,592 cells before and after stimulation with varying concentrations of DAP-type peptidoglycan (PGN) to activate the IMD pathway (Figure 1C). We developed an image processing pipeline to quantify nuclear and cytoplasmic Relish signals over time, then calculated the nuclear fraction, R_nuc:tot_, as an internally normalized measure of pathway activation for each cell ^30^ (Figure 1D).

In the absence of PGN, R_nuc:tot_ fluctuated within 20% of its initial value, which we used to define a threshold of pathway activation (Figure S2). Upon stimulation with 10 µg/mL of PGN, we observed an increase above threshold in the R_nuc:tot_ value of about 75% of cells. Despite the shared genetic background, we observed considerable cell-to-cell variability in both the timing and intensity of Relish nuclear translocation among these responders (Figure 1E).

### Relish dynamics follow a dose-dependent distribution of behaviors

Both the pre-stimulus and post-stimulus values of R_nuc:tot_ demonstrated heterogeneity across the cell population. We noticed that after stimulation, some cells seemed to rapidly translocate Relish to the nucleus, while others were slower. Similarly, some cells demonstrated a sustained nuclear localization of Relish over the 13 hour time course, while in others, R_nuc:tot_ diminished over time. To facilitate direct comparison of Relish nuclear-dynamics shapes independent of baseline, we normalized each full R_nuc:tot_ time trace by its mean pre-stimulus value, yielding the fold-change (FC) of R_nuc:tot_ over time. To characterize the diversity in these fold-change patterns, we manually classified a subset of traces based on their short-term and long-term behaviors (Figure S3). From this manual classification, five distinct categories of FC R_nuc:tot_ response dynamics emerged: Immediate (I), Immediate with Decay (Id), Gradual (G), Delayed (D), and Nonresponsive (N).

We converted these heuristic guidelines into a more rigorous classification framework by training a Support Vector Machine (SVM), hereafter called the Classifier SVM, to enable high-throughput categorization of single-cell FC R_nuc:tot_ traces. To build the SVM, we first extracted four quantitative descriptors that summarize dynamic properties of R_nuc:tot_ for each cell: the maximum FC value of R_nuc:tot_, the times to reach the maximum and half-maximum FC values, and the total area under the FC R_nuc:tot_ curve (Figure 2A). We manually labeled 150 cells using the heuristic guidelines above to train the SVM. 5-fold cross-validation results demonstrated an average classification accuracy of 87.3% on the testing set.

**Figure 2.**
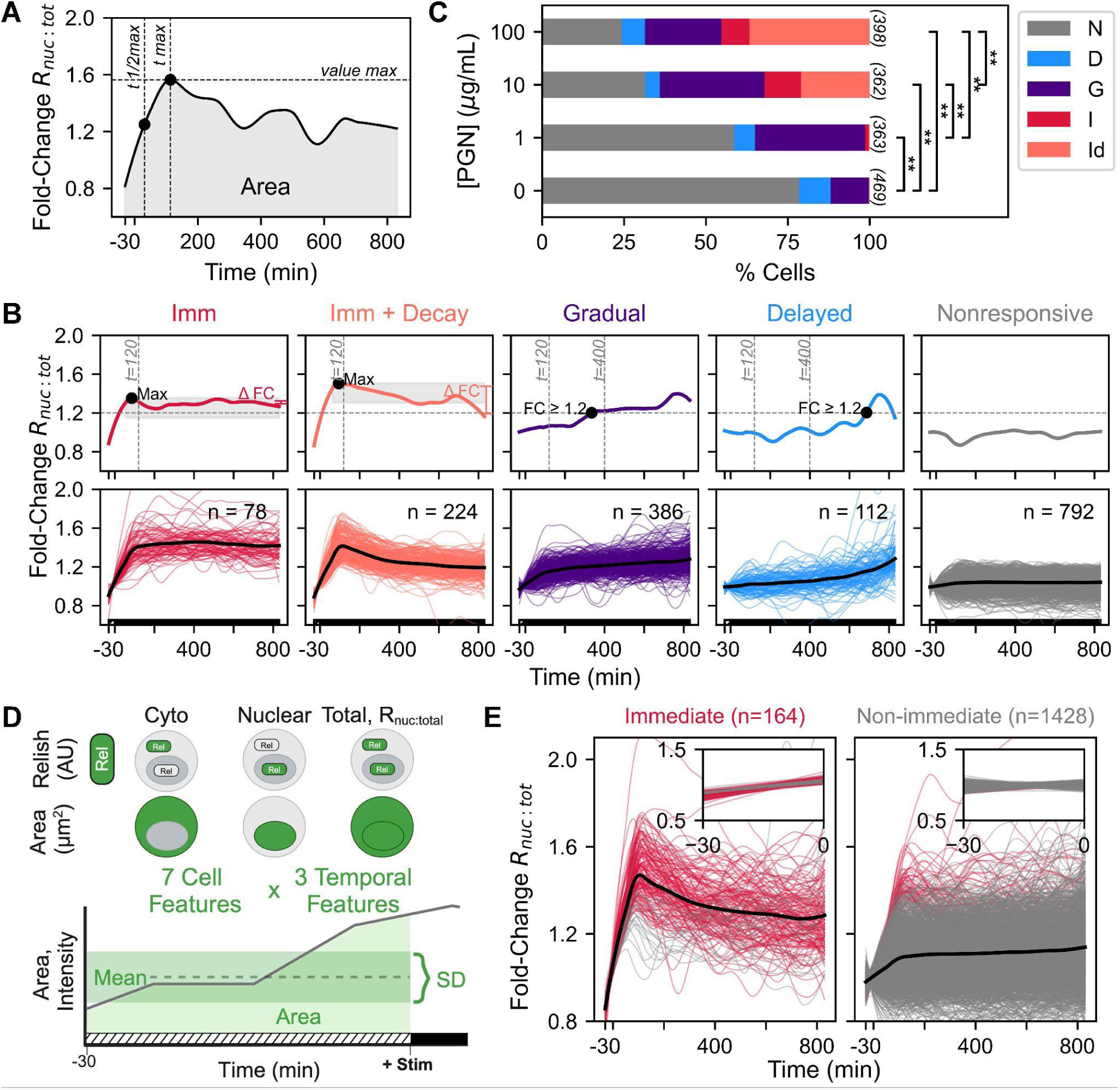
Classification of short- and long-term R_nuc:tot_ responses into five behavioral categories, with short-term responses predicted by pre-stimulus characteristics **A.** Schematic of the five extracted features of fold-change R_nuc:tot_ traces used to train the Classifier SVM: the maximum and half-maximum fold-change values, their associated times, and the area under the curve. **B.** The Classifier SVM distinguishes five Relish dynamics categories (top) and assigns each single-cell trace (bottom) (n = 1,592 cells from 4 independent replicates). Cells across 0, 1, 10, and 100 µg/mL PGN injection conditions are overlaid. Colored lines represent single-cell traces and overlaid bold line represents the mean of each category. **C.** Bar graphs depicting the percent of cells per Relish behavior category across PGN stimulation concentrations. Significance was determined by a χ² test with false discovery rate (FDR) correction (** p < 0.01). Corrected p-values for concentration pairs are included in Supplementary Table 2A. **D.** Schematic of the 21 pre-stimulus features used to train Predictor SVM. For each of the 7 features (absolute cytoplasmic, nuclear, and total Relish intensity per pixel, R_nuc:tot_, and cytoplasmic, nuclear, and total areas), the mean, SD, and AUC was calculated over the 30 minute pre-stimulus interval. **E.** The Predictor SVM categorizes cells as immediate (I, Id) or non-immediate (N, G, D) responders with 93.7% accuracy (n = 1592 cells). Red lines indicate single-cell traces of cells within I or Id categories, gray as cells within N,G, or D categories, as determined by the Classifier SVM. Overlaid bold line represents the mean of immediate and non-immediate category traces. Inset figure shows associated cells’ absolute R_nuc:tot_ magnified to pre-stimulus times. Horizontal bar on x-axis denotes the pre-(hatched) and post-(solid) injection phases.

To characterize the prevalence of the five behavior categories in our entire dataset of 1,592 cells, we used the SVM to classify each R_nuc:tot_ trace (Figure 2B). Examining behavior distributions across PGN stimulation concentrations reveals distinct patterns (Figure 2C). Fewer than 10% of cells exhibited Delayed behavior at any PGN concentration. As the PGN concentration increased, we observed a clear shift in behavioral responses: higher concentrations led to a greater proportion of Immediate and Immediate with Decay behaviors, while the ratio of Nonresponsive and Gradual cells decreased. Thus, increasing PGN concentrations increase both the proportion of responsive cells and the kinetics of their responses, consistent with a concentration-dependent progression likely reflecting saturation due to receptor occupancy limits. ^5,31^. In sum, our analysis reveals distinct Relish spatiotemporal behaviors across PGN-stimulated cells, with both responsiveness and response speed increasing with concentration.

### Pre-stimulus Relish subcellular localization predicts pathway responsiveness

Unsurprisingly, we observed a non-uniform baseline level of nuclear Relish across the cell population prior to IMD induction (Figure S4). We hypothesized that each cell’s pre-stimulus state could influence its behavioral outcome, and to test this hypothesis, we developed another SVM model (referred to as Predictor SVM) trained on 21 parameters derived from seven cellular pre-stimulus features: R_nuc:tot_; average cytoplasmic, nuclear, and total Relish intensity; and cytoplasmic, nuclear, and total areas. In the 30 minute pre-stimulus interval, we imaged the cells every 10 minutes, yielding 4 images per cell. Therefore, for each feature, we calculated three metrics—mean, standard deviation, and area under the curve, providing a comprehensive representation of the cell’s pre-stimulus state (Figure 2D, Figure S5). While cells may change states over longer time scales, a previous study in mammalian cells demonstrated that ∼30 minutes is a sufficient sampling duration immediately prior to stimulation to capture a cell’s potential to activate ^9^.

We reasoned that cells’ immediate post-stimulus response would more robustly be predicted by their pre-stimulus state, compared to their emerging long-term behavior. Therefore, in this version of the Predictor SVM, we labeled cells in both the Immediate and Immediate with Decay categories as “immediate” and the Gradual, Delayed, and Non-responsive categories as “non-immediate.” Our model’s prediction accuracy was 93.7% (Figure 2E), indicating that the pre-stimulus Relish and area parameters can robustly distinguish between cells that will immediately respond to stimulus and those that will not. Upon closer investigation of the pre-stimulus absolute R_nuc:tot_ values (insets, Figure 2E), the slopes of immediate responders’ R_nuc:tot_ trajectories appear higher than those of non-immediate responders, indicating that cells with an already increasing R_nuc:tot_ ratio may be primed to respond.

To pinpoint the most informative among the 21 features—some of which were temporally correlated—we applied recursive feature elimination. This analysis showed that a core set of three metrics (the standard deviation of the Relish nuclear fraction, the AUC of the Relish nuclear fraction, and the AUC of nuclear Relish) was sufficient to sustain strong classification performance, with the reduced model reaching 89.4% accuracy (Figure S5).

To test whether pre-stimulus parameters could be used to predict behavior beyond two hours post-stimulus, we created modified versions of the Predictor SVM. The first modified version attempted to classify cells’ full dynamic behavior (I, Id, G, D, or N); the second version classified behaviors collapsed into either nonresponsive vs. responsive (N vs. [I, Id, G, D]); the third version sorted into nonresponsive, immediate, and long-term categories (N vs. [I, Id] vs. [G, D]). These modified Predictor SVMs achieved slightly lower but still robust accuracy rates of 76.9%, 81.5%, and 78.6%, respectively. This indicates that while a model using pre-stimulus cell characteristics may accurately predict long-term dynamics, it is best at predicting behavior within two hours after exposure to stimulus.

### Simultaneous tracking of Relish dynamics and AMP transcription reveals Relish-dependent expression patterns

Upon immune stimulation, nuclear Relish activates the expression of various AMPs that are secreted to combat invading pathogens. To measure how Relish’s spatiotemporal heterogeneity may impact AMP transcription, we built transcriptional reporter cell lines to simultaneously track these two nodes of the pathway. AMPs of interest included the IMD-responsive Diptericin (Dpt) and IMD and Toll-responsive Metchnikowin (Mtk) ^21,32–34^, both of which display fast and strong upregulation within 1–2 hours after infection in live flies ^32^. IMD-responsive enhancer regions were identified for Dpt and Mtk with a large-scale reporter assay (STARR-Seq) conducted in S2* cells ^35^ 24 hours after stimulation by heat-killed *Serratia marcescens*, which presents IMD-stimulating DAP-type PGN. The identified regulatory region for Dpt is located between the *DptA* and *DptB* genes, partially overlapping with *DptB*, whereas Mtk’s regulatory region is situated within the *Mtk* gene itself (Supplementary Figure S6). We additionally designed a reporter for the Toll-responsive AMP, BomS2 (also known as IM2), as a pathway-specific negative control. No enhancer was identified for BomS2 in the IMD-stimulated reporter assay, so we designed the enhancer reporter to include two Toll-responsive κB motifs upstream of the gene ^34^. This negative control reporter should not display transcription in response to PGN, as prior studies found that the *BomS2* gene is predominantly activated in response to Lys-type PGN ^33^, and the included κB motifs are more strongly bound by the Toll-responsive NF-κB, DIF34_._

To track transcriptional dynamics, we modified the previous cassette to include the enhancer sequences upstream of RhoBAST RNA aptamer repeats ^24^, flanking the assembly with insulating sequences to sequester enhancer activity from the constitutive promoter driving Relish expression ^36^ (Figure 3A, S6). When transcribed, the RhoBAST sequence folds into mRNA stem loop structures that can be labeled by rhodamine dyes, such as SpyRHO555 ^24^. As transcriptional dynamics are governed by both slow processes, such as nucleosome remodeling, and fast processes, such as the formation and disassembly of transcriptional complexes ^37^, we imaged the reporter lines at two different time scales to capture both the long-term transcriptional persistence and the temporally-resolved transcriptional burst dynamics. A sparse imaging setup (one frame per 15, 30, then 60 minutes) followed the same fields of view longitudinally and minimized cellular photobleaching, whereas a dense imaging setup (two frames per minute) captured different cells within three evenly-spaced windows of time across the full duration (Figure 3B).

**Figure 3.**
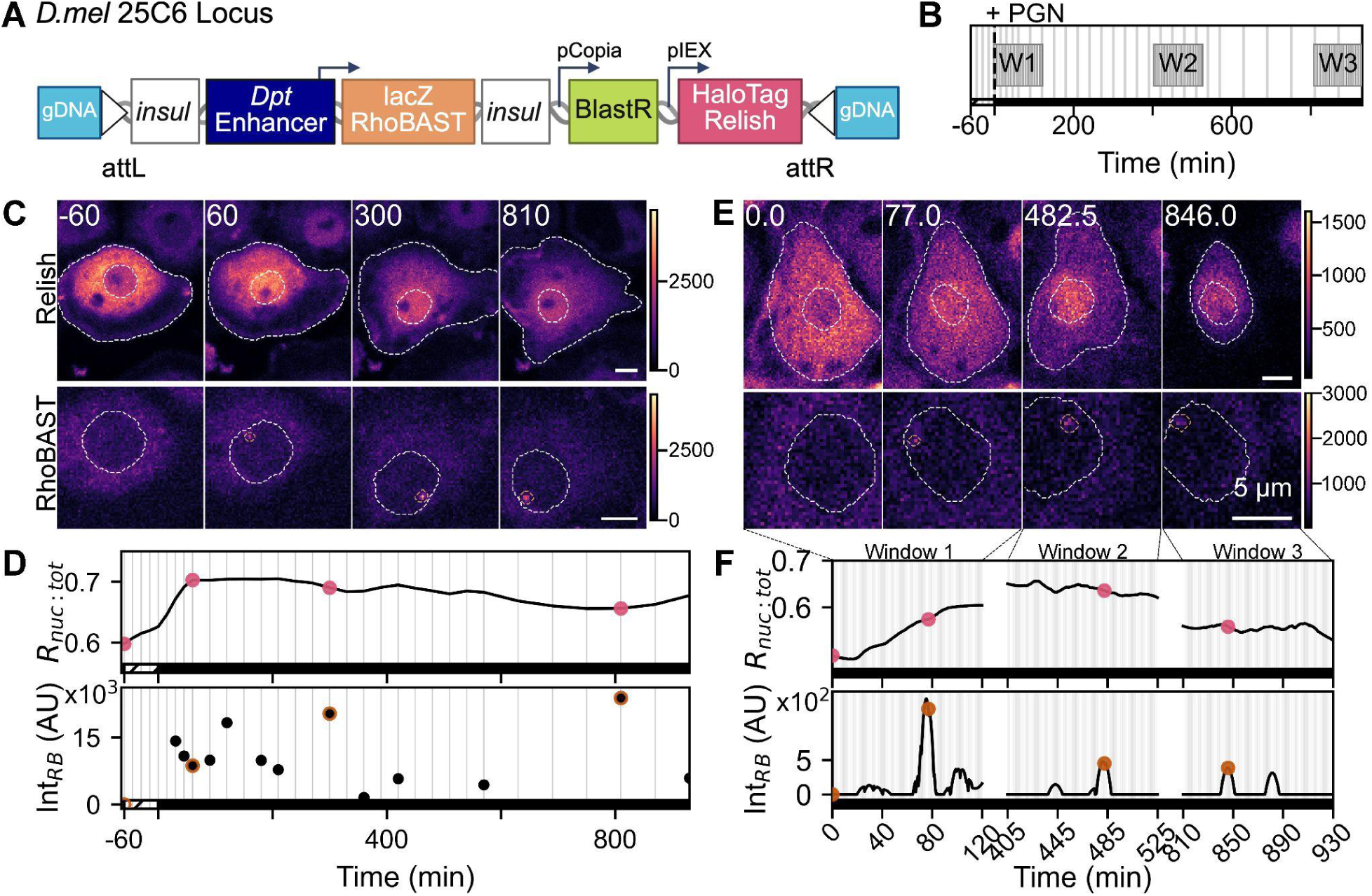
Dual imaging setups for simultaneous analysis of AMP enhancer-driven transcription and Relish dynamics. **A.** Schematic illustrating transcriptional reporter inserted via RMCE at attP sites ^29^. AMP enhancer (with encompassing promoter region) drives the transcription of lacZ-8xbiRhoBAST aptamer, with enhancer activity insulated from downstream Relish reporter. **B.** Schematic of dual imaging modalities. Sparse acquisition collected frames every 15 minutes for 2 hours (with 1 hour pre-stimulus baseline), every 30 minutes for 8 hours, then every hour for 5 hours (interspersed light grey lines). Dense acquisition collected frames immediately following stimulus, every 30 seconds in two-hour windows (labeled W1, 2, and 3). **C.** A representative cell imaged with the sparse setup, depicting the signal of HaloTag-labeled Relish (top) and SpyRHO555-labeled RhoBAST aptamer (bottom), both pre-stimulus (t = -60), and post-stimulus with 100 µg/mL PGN (t = 60, 300, 810 minutes). Overlaid masks show cell bodies and nuclei (top panels), and the same nuclei containing segmented RhoBAST foci (bottom panels). Scale bar = 5 µm. Images are confocal maximum-intensity projections (three z-steps spanning 2 µm). **D.** Quantification of R_nuc:tot_ (top) and sum of segmented RhoBAST intensity (bottom) of cell in C over time. R_nuc:tot_ was smoothed over time using Savitzky Golay filtering with a polynomial order of 2. Marked timepoints in pink and orange indicate selected Relish and RhoBAST panels in C, respectively. **E.** Representative cell imaged with dense configuration with panels organized as in C. Cell was imaged post-stimulus with 100 µg/mL PGN with panels selected from window 1 (t = 0.0 and 77 seconds), window 2 (t = 482.5 seconds), and window 3 (t = 846 seconds). Images are a single confocal z-slice. Scale bar = 5 µm. **F.** Quantification of R_nuc:tot_ (top) and sum of segmented RhoBAST intensity (Int_RB_, bottom) of cell in E over time. RhoBAST foci traces were considered continuous with imaging every 30 seconds. R_nuc:tot_ and Int_RB_ were smoothed over time using Savitzky Golay filtering with a polynomial order of 2. Marked timepoints in pink and orange indicate selected Relish and RhoBAST panels in E, respectively.

Our complementary imaging approaches revealed a trade-off in acquisition strategy, with sparse imaging effectively capturing cells’ respective R_nuc:tot_ dynamics but missing individual bursts, while dense imaging successfully resolved burst dynamics but prevented long-term tracking of nuclear Relish localization. Following PGN injection, sparse imaging of the *Dpt* enhancer reporter line revealed prominent foci of RhoBAST aptamer transcription within the nuclei of cells also displaying nuclear-localized Relish (Figure 3C, Supplementary Video 1). In contrast, imaging of the negative control *BomS2* reporter, or any enhancer in the absence of PGN, produced very few detectable foci (Figure 4A, Figure S7B). As in previous studies ^38–41^, we interpret these bright foci to be the site of nascent transcription and conclude that our imaging approach allows us to visualize clusters of mRNA molecules but is not sensitive enough to detect individual mRNA molecules farther from the site of transcription. To estimate the nascent RhoBAST RNA abundance, we segmented the nuclear foci and summed their total intensity over pixels (Int_RB_) across time frames. As seen in the sample trace exhibiting Immediate Relish behavior (Figure 3D, top), this approach allowed us to capture the cell’s full R_nuc:tot_ spatiotemporal behavior as before, but likely missed transcriptional events occurring between frames (Figure 3D, bottom). In contrast, the dense imaging setup captured complete RhoBAST bursts (Figure 3E, 3F, bottom, Supplementary Video 2), enabling quantification of transcriptional features such as burst duration and slope as a function of AMP enhancer. However, the high sampling rate required to capture these bursts resulted in cell photobleaching and phototoxicity of nearly all cells, preventing longitudinal tracking of the cells’ R_nuc:tot_ behavior over the full duration of imaging (Figure 3E, top). We therefore chose to analyze different cells across each time window to optimize cell health in this dataset. We use both imaging approaches below to probe different questions about Relish-driven transcription.

**Figure 4.**
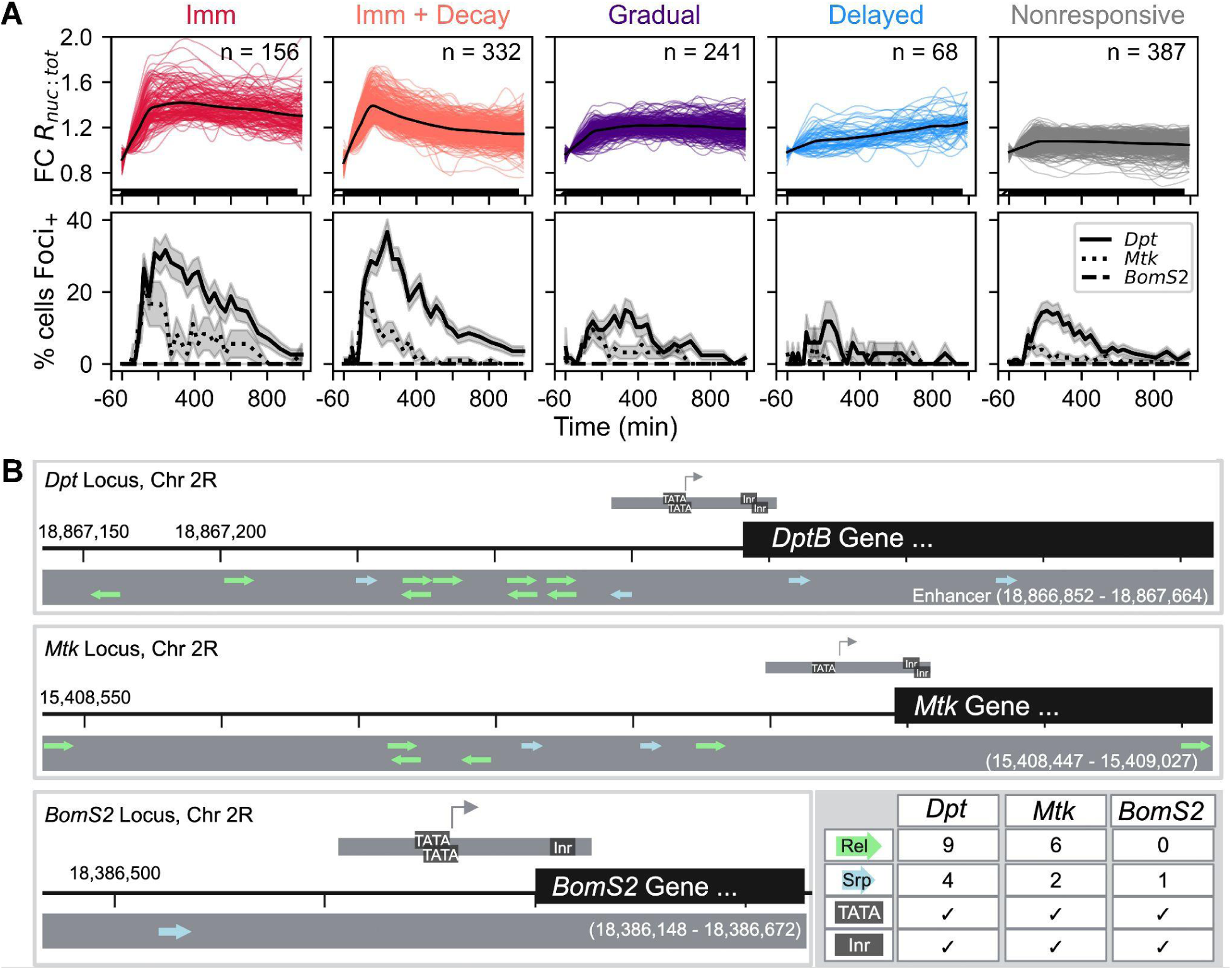
Impact of Relish spatiotemporal behavior and enhancer binding site number on ratio of transcriptionally active cells. **A.** Transcriptional reporter cells stimulated with 100 µg/mL PGN were categorized by the Classifier SVM (top, n = 1,184 cells from 4 independent replicates). Colored lines represent single-cell traces and overlaid bold line represents the mean of each category. Of the cells within each category, the percent displaying spots of transcriptional foci at each time point is plotted across reporters for Dpt, Mtk, and BomS2 AMPs (bottom). Traces represent the mean ± SEM of all cells within categories **B.** Enhancer sequences for *Dpt*, *Mtk*, and *BomS2* were scanned for core promoter TATA box and initiator (Inr) sequences, as well as for binding site motifs for the transcription factors Relish and Serpent. Each contained a similar promoter architecture but variable numbers of binding sites for Relish and Serpent.

To understand how different enhancer reporters respond to Relish activation, we once again categorized the single-cell Relish dynamics into five behavior categories following stimulation with 100 µg/mL of PGN, then measured the concurrent percentage of transcriptionally active reporters over the time course (Figure 4A, top). Overall, transcriptional activity (measured by the presence of RhoBAST foci) varied both across Relish behavior categories and AMP enhancer (Figure 4A, bottom). Across all Relish behaviors, *Dpt* reporters consistently showed the highest proportion of transcriptionally active cells, followed by *Mtk*, with *BomS2* exhibiting minimal activity, as expected. For example, in the Immediate categories, a peak of ∼35% *Dpt* reporter cells had a RhoBAST focus, while *Mtk* reporters in these categories peaked at ∼21% of cells transcribing.

The temporal transcriptional dynamics of *Dpt* and *Mtk* reporters broadly mirrored the corresponding Relish behavior, with slight reporter-specific differences. For example, cells in the Gradual Relish category showed delayed transcriptional activity compared to the Immediate categories, and cells in the Immediate category showed sustained transcriptional activity compared to those in the Immediate with Decay category. Neither enhancer displayed much transcriptional activity when Relish behavior fell into the Delayed category, suggesting that this pattern may not be indicative of strong IMD stimulation. Similar dynamics were observed across cells stimulated with 10 µg/mL PGN, although at generally lower percentages of responding cells (Figure S7). A small ratio of *Dpt* and *Mtk* reporter cells exhibits foci in the Nonresponsive category, which indicates that some cells that are below the FC R_nuc:tot_ 20% activation threshold are still capable of initiating reporter transcription.

The timing and duration of *Dpt* and *Mtk* reporter activity differed in consistent, reporter-specific ways across Relish behavior categories. In the Immediate Relish category, nuclear Relish peaked at around 200 minutes post-stimulus, with the *Dpt* reporter peaking 40 minutes later and the *Mtk* reporter peaking 80 minutes earlier. Similar temporal offsets were observed in the Immediate with Decay category, where nuclear Relish peaked at 166 minutes, followed by *Dpt* peaking 70 minutes later and *Mtk* peaking 46 minutes earlier. Notably, *Mtk* reporter activity consistently peaked earlier and decayed more rapidly than *Dpt* across Relish behavior categories, and both reporters’ activities declined more quickly than nuclear Relish levels. At a broad level, these data demonstrate that transcriptional activity is correlated with upstream Relish dynamics, as expected, but reveal quantitative differences in both relative timing and overall persistence of activity between the two Relish-responsive reporters.

### Transcription factor binding site composition underlies differential enhancer activity

To investigate the source of varying transcriptional output across the reporters, we compared the regulatory motifs present within the AMP enhancers (Figure 4B). Using the Elements Navigation Tool ^42^, we searched for the *Drosophila* core promoter elements that recruit the core promoter recognition factor (TFIID) responsible for assembling the pre-initiation complex driving transcription. These elements include motifs such as the TATA box, which binds the TATA-binding protein (TBP) upstream of the transcriptional start site, and downstream promoter sequences such as the initiator element (Inr), which binds TBP-associated factors ^37^. We found both TATA boxes and Inr motifs within the previously established promoter regions of each AMP ^43^, and while the number of elements varied across enhancers, they collectively formed a similar promoter configuration, maintaining the core regulatory components necessary for transcription initiation (Figure 4B, grey boxes above gene).

Given the similar promoter architectures, we next investigated whether the number of transcription factor binding sites could explain the observed transcriptional variation across enhancers. Using the MEME suite’s FIMO tool ^44,45^, we scanned for motifs for various immune-responsive transcription factors, and identified binding sites for Relish and Serpent, a GATA factor required for hemocyte development^46^. Notably, we found the highest number of binding sites in the *Dpt* enhancer, including nine for Relish and four for Serpent. In comparison, the *Mtk* enhancer contained six Relish and two Serpent sites, while the *BomS2* enhancer only contained one Serpent site (Figure 4B, blue and green arrows). Serpent is a hemocyte lineage-defining transcription factor that cannot induce immune-responsive transcription on its own, but previous work has shown that Relish and Serpent sites can work synergistically to activate AMP transcription ^47^. These findings align with our data, which show *Dpt* cells as the most transcriptionally active and *BomS2* cells as the least across all Relish behaviors. This suggests that the number of binding sites in each enhancer is a key factor in determining the strength of AMP activation following immune stimulation.

### Instantaneous Relish nuclear fraction predicts AMP transcriptional activity

To further explore the relationship between Relish spatiotemporal dynamics and transcriptional activity, we investigated which features were most predictive of the occurrence of transcriptional foci from the *Dpt* and *Mtk* enhancer reporters. We assigned each of the 28 post-stimulus movie frames of the sparsely imaged cells as either tFoci+ or tFoci-, determined by whether or not a RhoBAST peak was detected in that particular frame (Figure 5A). For *Dpt*, this resulted in 2025 tFoci+ frames and 16,259 tFoci-frames across 653 cells, and for *Mtk*, 387 tFoci+ frames and 12,129 tFoci-frames across 447 cells (Figure 5B). We first tested how predictive the concurrent R_nuc:tot_ value – the value from the same movie frame – would be at discriminating between tFoci+ and tFoci-frames by plotting both the raw Relish nuclear fraction (R_nuc:tot_) and pre-stimulus value-normalized fraction (FC R_nuc:tot_) distributions (Figure 5C). Distributions of both normalized and non-normalized R_nuc:tot_ values for tFoci+ and tFoci-frames proved to be significantly different for both *Dpt* and *Mtk* reporter cells, suggesting that the concurrent Relish behavior was predictive of transcription occurrence. The corresponding Receiver Operating Characteristic (ROC) curves for the distributions further characterized the performance of this binary classification across all thresholds of R_nuc:tot_: the unnormalized R_nuc:tot_ values for each time point proved to be significantly more predictive of transcription than the corresponding normalized value, as indicated by the larger area under the curve for both *Dpt* and *Mtk* (0.73 vs. 0.70 for *Dpt*, 0.77 vs 0.71 for *Mtk*, respectively).

**Figure 5.**
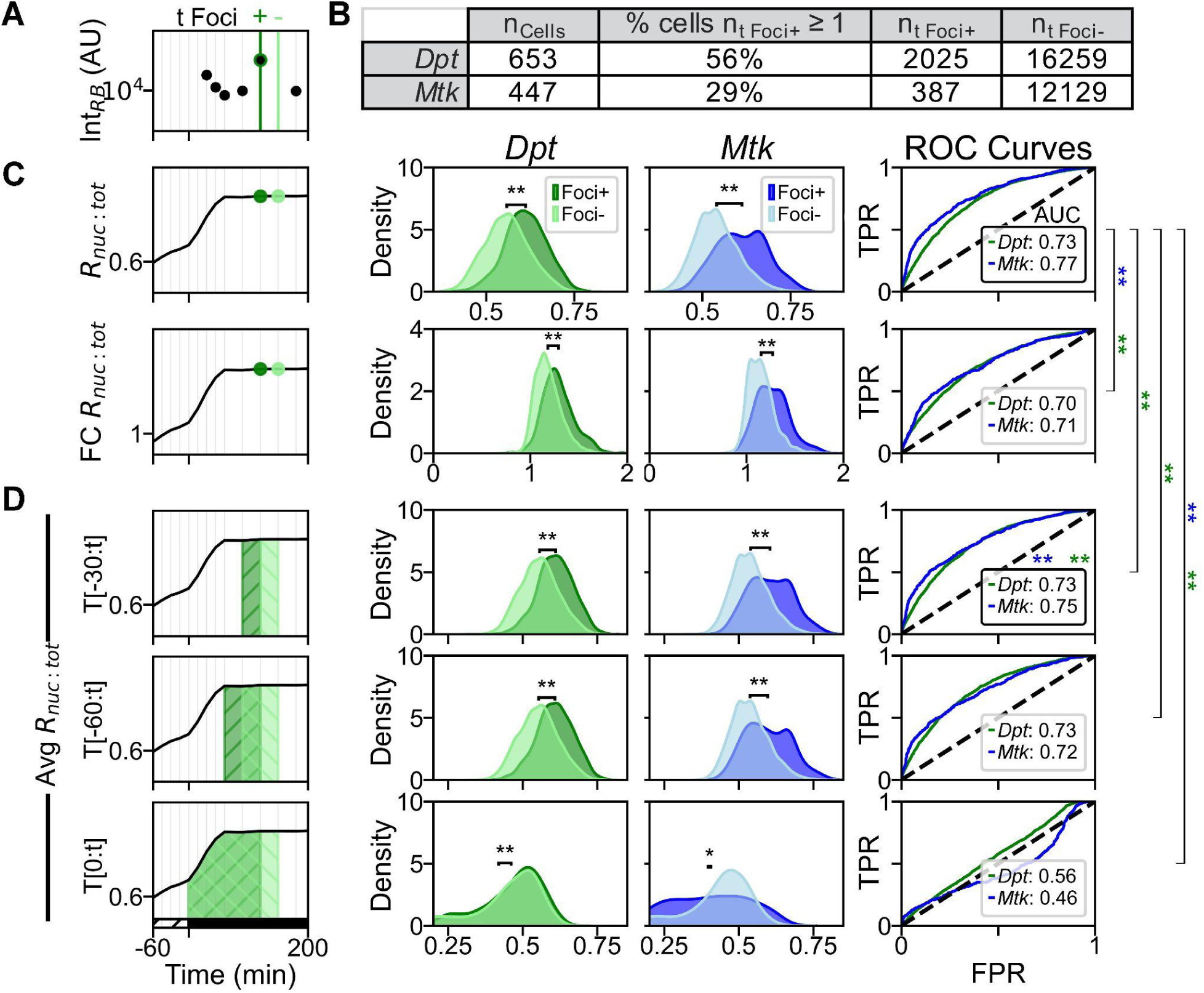
Prediction of transcriptional events by short-term R_nuc:tot_ levels. **A.** Schematic illustrating classification of sparse-imaging time frames as transcriptionally active (tFoci+) or inactive (tFoci-) based on presence of nuclear RhoBAST foci. **B.** Summary table of cell counts, percentage of cells with at least one tFoci+ frame, and total timepoints with or without foci for *Dpt* and *Mtk* enhancer reporters. **C-D.** Quantitative analysis of Relish nuclear localization features and their predictive power for AMP transcription. *Left*: Schematic representations of the Relish nuclear localization metrics evaluated, including instantaneous R_nuc:tot_, instantaneous fold-change (FC) R_nuc:tot_ **(C)**, and time-averaged R_nuc:tot_ over different windows relative to t = tFoci+ (T[-30:t], T[-60:t], T[0:t]) **(D)**. *Middle*: Kernel density estimates of each metric for Foci+ (dark) and Foci– (light) frames, shown separately for *Dpt* (green) and *Mtk* (blue) reporters. Significance was determined by a Kolmogorov–Smirnov test (* p < 0.05, ** p < 0.01). P-values for *Dpt* and *Mtk* tFoci+ vs tFoci-distributions across each metric are included in Supplementary Tables 2B. *Right*: Receiver Operating Characteristic (ROC) curves and AUC values for each curve quantify the ability of each metric to discriminate between transcriptionally active and inactive frames. Instantaneous R_nuc:tot_ values were most predictive of all metrics. Differences in AUC values across metrics was determined by a DeLong’s test with Bonferroni correction (** p < 0.01). Corrected p-values for metric pairs are included in Supplementary Tables 2B.

Despite the predictive ability of a single frame’s Relish behavior, this metric reveals nothing about the cell’s complex past signaling history. It is possible for NF-κB signaling and target gene expression to show discordant levels at snapshot readouts due to the lag of activation and accumulation of target transcripts ^18^. For example, we may observe a high R_nuc:tot_ value without a transcriptional focus if Relish has just entered the nucleus and there is a lag prior to transcriptional activation. We therefore quantified the time-integrated activity of Relish across different time windows by calculating the area under the R_nuc:tot_ plots leading up to each frame (Figure 5D). We tested the predictive power of normalized AUC, which corresponds to the average R_nuc:tot_, from the beginning of stimulus (t = 0), 60 minutes (t = -60), and 30 minutes (t = -30) leading up to the tFoci+ or tFoci-frame by normalizing the integral to the duration of the integration window. Interestingly, the distributions of tFoci+ and tFoci-frames were significantly different irrespective of the time of integration for both *Dpt* and *Mtk* reporters. Across both enhancers, the area under the ROC curves indicated that the t = 0 curve performed significantly worse than t = -60 and t = -30, and that the t = -30 curve was significantly more predictive than the longer integrations. From these analyses, we find that the raw R_nuc:tot_ value is more predictive of the presence of a transcriptional burst than the FC value, in contrast to what has been found in some mammalian NF-κB systems ^48^. Despite the caveats of the time lags in some measurements, the ROC AUC metric can exceed 0.7, indicating that the transcriptional behavior of our reporters is well-explained by this metric. Further, integrating R_nuc:tot_ to account for prior signaling behavior does not improve predictive power, suggesting that the transcriptional signaling is not dependent on the prior Relish nuclear fraction.

### Transcriptional probability and initiation rate vary between enhancers

While the sparse imaging configuration enabled us to explore how enhancer activity correlates with long-term Relish dynamics, we also aimed to capture transcriptional burst dynamics between enhancers at a finer temporal resolution. The dense imaging setup allowed us to achieve this higher resolution of transcriptional bursts (Figure 3B). Our reporter starts with the lacZ sequence, followed by a 3’ RhoBAST aptamer, which means transcription starts prior to labeling with SpyRHO555. To account for this, we shifted all peak start times two minutes earlier—reflecting the approximate duration needed to transcribe the lacZ region, assuming a transcription rate of about 1.5 kb/minute ^41^ (Figure 6A, inset).

**Figure 6.**
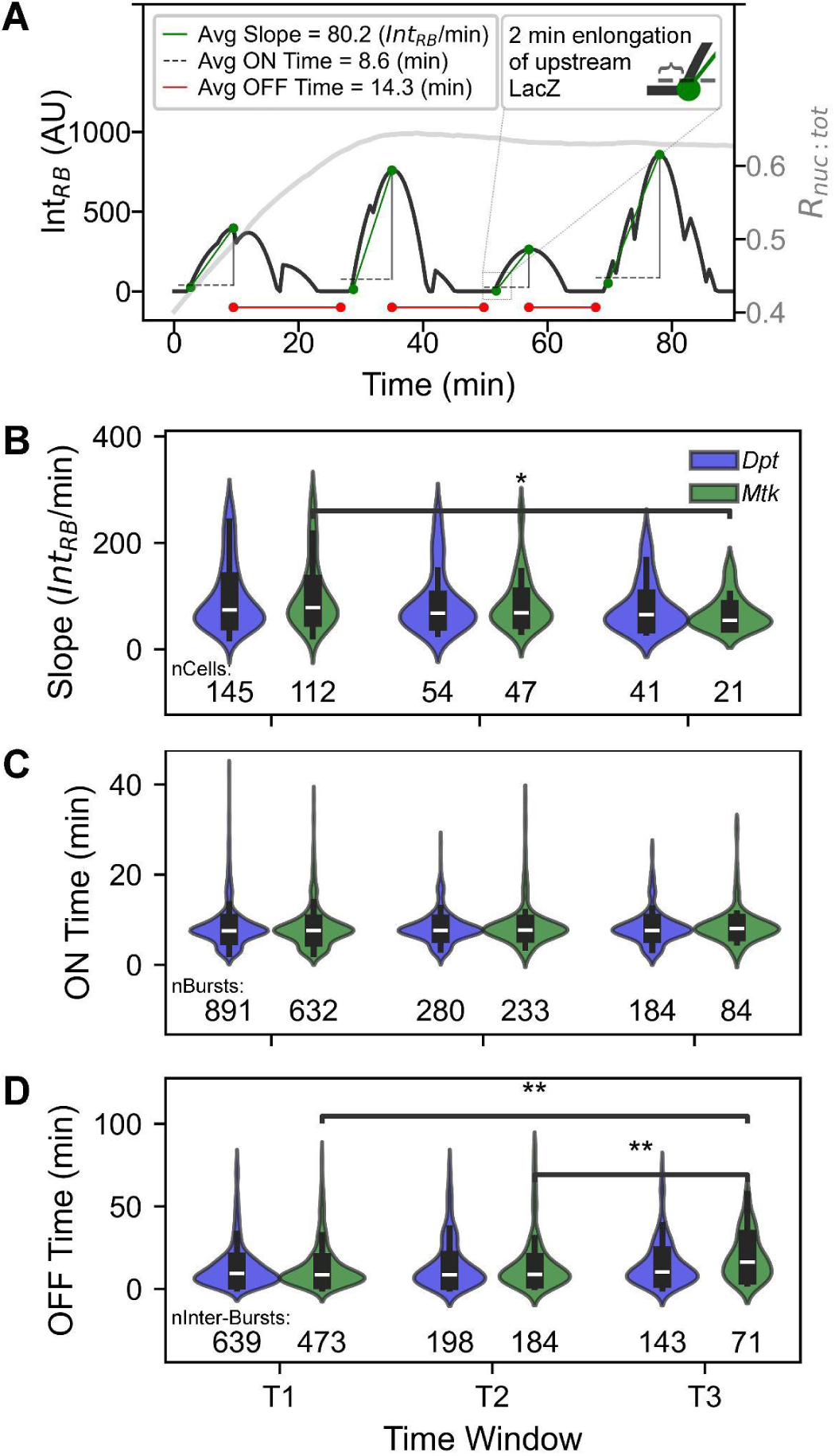
*Mtk* transcriptional burst slopes and OFF times, but not ON times, vary across time windows. **A.** Example *Dpt* trace of total RhoBAST fluorescence (Int_RB_, left y-axis) and R_nuc:tot_ (right y-axis) during time window 1, and the measured burst characteristics: average slope of fluorescence increase during a burst (green line, 80.2 AU/min), average burst ON duration (dashed line, 8.6 min), and average burst OFF duration (red line, 14.3 min). Inset: burst start time is adjusted to include elongation time of 5’ lacZ sequence. **B-D.** Comparison of burst parameters for *Dpt* (blue) and *Mtk* (green) reporter cells across three dense imaging time windows: **(B)** Average slope of fluorescence increase for each bursting cell (top 90% of observed bursts plotted, Int_RB_/min), **(C)** Burst ON durations, and **(D)** Burst OFF durations for individual bursts. Violin plots show the distributions for each group (median ± IQR as boxplots), with n values indicated below each category (data collected over 6 independent replicates). Significance was determined by a two-sided Mann–Whitney test with Bonferroni correction (* p < 0.05, ** p < 0.01). Corrected p-values for significant pairs are included in Supplementary Table 2C.

For each cell with detected RhoBAST transcriptional bursts, we measured the average slope of bursts during periods of active transcription, which approximates the rate at which RNA polymerase loads and actively transcribes the reporter (Figure 6A, B). The average slope of *Dpt*-enhancer-driven bursts was 139.6 ± 150.1 units/minute, with no significant difference across the three time windows. *Mtk* reporters initially displayed a slope of 151 units/minute, which then decreased significantly to 82.1 units/minute by window 3 (Figure 6B). By adjusting transcription rate without altering burst duration (Figure 6C), the *Mtk* enhancer modulates burst size. This is consistent with previous findings that highly expressed loci increase activity by modulating burst size in response to immune stimulation ^49,50^. These data indicate that the *Mtk* enhancer attenuates activity over time after stimulus by reducing RNA polymerase loading rate, and therefore burst size, whereas *Dpt* maintains a steady, high rate of transcription along the same time course.

We divided transcribing cells’ RhoBAST traces into periods of active transcription (ON) and intervening (OFF) intervals to quantify each enhancer’s characteristic burst ON and OFF times (Figure 6A). We found no significant change in burst ON times across enhancers or time windows, with an average of 8.2 ± 4.2 minutes across all conditions (Figure 6C). While *Dpt* reporters’ burst OFF times remained at 13.7 ± 13.5 minutes across all time windows, *Mtk* reporters’ OFF times significantly increased from 13 ± 13.1 to 14.6 ± 15.6 minutes from window 1 to 2, and further to 19.4 ± 13.8 minutes by window 3. We next investigated whether the immediacy of Relish nuclear localization impacted the ON or OFF time distributions, but found no significant effect (Figure S8B-D). The *Mtk* enhancer thus becomes less active over time following IMD stimulus by both attenuating burst size and increasing the waiting time between bursts, whereas the *Dpt* enhancer drives consistent-sized bursts at regular intervals over the course of 15 hours post-stimulus.

Transcriptional bursting can be represented by a two-state promoter model in which a gene stochastically switches between ON and OFF states. The ON and OFF times are assumed to be regulated by single rate-limiting steps, with exponentially distributed time between bursts ^51^. However, across most time windows, burst ON and OFF durations in our data did not fit well to an exponential distribution (Figure S8A). Burst ON and OFF times were better described with a non-monotonic unimodal distribution. In previous modeling work aimed to adapt the two-state promoter model to recapitulate non-exponential waiting times, researchers found that adding a third refractory state to enforce a minimum “off” time between bursts or incorporating oscillatory modulation of gene activation were both possible mechanisms ^37,52^.

Collectively, characterization of transcriptional burst dynamics suggests different mechanisms of temporal regulation between the two enhancers following IMD stimulation. Both enhancers maintain stable durations of active transcription once initiated, but *Dpt* transcription speed and frequency do not decrease as quickly at *Mtk* over time. Furthermore, the consistent ON times of *Dpt* and *Mtk* bursts over time suggests we are unlikely to be selectively missing foci from one reporter over the other in our sparse imaging. Therefore, the ratio of responding cells in the population is likely also attenuated over time, more so with *Mtk* than *Dpt* (Figure 4A, bottom). Together, these results indicate that *Mtk* cells respond more quickly to stimulation than *Dpt*, but drive an overall lower amount of AMP expression due to its faster attenuation of transcription rate and ratio of transcriptionally active cells within the population.

## Discussion

Innate immunity is a vital, evolutionarily conserved defense against pathogenic infections, yet how it is regulated at the single-cell level remains incompletely understood. The diverse responses of individual cells within an immune-responsive population can influence the overall magnitude, timing, and coordination of the organismal response to infection, highlighting the need to uncover how such variability arises. In this work, we engineered novel live-cell reporters to identify sources of signaling heterogeneity across the IMD innate immune pathway. We find heterogeneity both in the behaviors of NF-κB localization dynamics and in the transcriptional activity of cells within behavior categories, indicating variability at multiple layers of the pathway. By tracking the NF-κB transcription factor, Relish, across a large population of cells, we find distinct categories of spatiotemporal behavior following pathway stimulation with PGN, which we refer to as Immediate, Immediate with Decay, Gradual, Delayed, and Nonresponsive (Figure 2B). We demonstrate that higher PGN concentrations increase not only the ratio of responsive cells, but also the fraction of cells in the Immediate categories (Figure 2C). A predictive SVM trained on 30 minutes of pre-stimulus data uncovers that a cell’s likelihood of rapid Relish activation is encoded in a few measurable parameters, most importantly its absolute nuclear Relish fraction (Figure 2D, E). We engineered our Relish reporter to include RNA aptamers driven by the *Diptericin* (*Dpt*), *Metchnikowi*n (*Mtk*), and *BomS2* enhancers, creating transcriptional reporter lines to simultaneously monitor each AMP enhancer’s transcriptional activity alongside upstream Relish dynamics. Expression was tightly but non-linearly coupled to upstream signaling, as AMP transcription followed promptly after Relish nuclear translocation, and the ratio of transcriptionally active cells was tuned by the number of κB binding sites within the enhancer (Figure 4A). The immediate nuclear Relish fraction strongly predicts transcriptional events, with no improvement when preceding Relish dynamics are averaged over time (Figure 5). Finally, high-frequency imaging of our transcriptional reporter cell lines uncovered AMP enhancer-specific modulation of burst slope (polymerase loading rate) and inter-burst intervals (OFF times) over time (Figure 6).

### Heterogeneity in Relish’s predisposition to activation, long-term dynamics, and downstream transcription

Despite the innate immune system’s ability to mount specific responses to varying stimuli concentrations, studies across organisms report vast heterogeneity among immune-responsive cells ^9,19–21,53^. We provide evidence that heterogeneity exists in cell populations even prior to immune stimulation, and that pre-stimulus states can be used to predict responsiveness (Figure 2D; Figure S5). Strikingly, predisposition to activation seems to be conserved across species, as similar models in mammalian immune systems could predict cellular responsiveness using pre-stimulus states as input ^9^. This idea is also supported at the organism scale by transcriptomic studies in *Drosophila* hemocytes. Single-cell RNA sequencing of unchallenged larvae uncovered distinct transcriptional clusters within the extracted hemocytes, with some clusters showing elevated Relish and AMP expression, suggesting the existence of cell reservoirs primed for humoral immune response ^19^.

In addition to heterogeneity in pre-stimulus states, immune challenge introduces another layer of variability across cell populations. At the NF-κB level, this was evident in our data as longitudinal Relish dynamics diverged into different spatiotemporal categories following PGN injection (Figure 2B, 4A). Notably, while our Predictor SVM model could predict immediate Relish behavior based on pre-stimulus states, it was less effective at forecasting long-term dynamics, suggesting that other mechanisms contribute to stochasticity in the long-term Relish behavior. At the transcription level, we observed heterogeneity in AMP enhancer dynamics (Figure 4A). Transcriptional heterogeneity has also been observed at the tissue level. For example, single-cell RNA sequencing of fly hemocytes revealed only a small subset of cells expressed AMPs in response to immune challenge ^20^. A single-cell RNA-seq study of the fly fat body, a highly immune-responsive organ similar to the mammalian liver, identified different cell clusters by their transcriptional signature. Nearly all cell clusters upregulated immune-responsive genes following infection, though particular gene combinations were more strongly induced in some clusters than others ^21^. These data demonstrate that the NF-κB response is heterogeneous, both pre- and post-stimulation and at the levels of NF-κB localization and ensuing transcriptional dynamics.

### Mechanisms driving heterogeneity

Subtle variations in each cell’s pre-existing regulatory landscape can give rise to heterogeneity within immune-responsive cell populations. Leading hypotheses for the factors driving these differences include cell cycle stage ^54^ and pre-existing epigenetic states such as chromatin accessibility ^50^ and levels of pathway components ^9,25,50,54,55^. In support of epigenetic factors’ influence on NF-κB response, daughter mouse fibroblast cells inherit some aspects of NF-κB response profile from their parent cell, as evidenced by their similar NF-κB oscillation dynamics ^54^. Studies also highlight that varying levels of pathway regulators can contribute to population heterogeneity. In mammalian systems, a high pre-stimulus protein ratio of NF-κB to its inhibitor, IκB, predicts leaky or rapid pathway activation ^9^. Analogously, the immune responsiveness of *Drosophila* cells may be influenced by the ratio of Relish with its negative regulators, Pirk and Charon ^2,56,57^. Investigating these mechanisms may uncover conserved principles of immune priming and the molecular logic of immune readiness. Collectively, these results underscore how random initial conditions or molecular fluctuations can tip one cell into full NF-κB activation, while a seemingly identical one in the population remains inactive under the same stimulus conditions. While much of this “noise” may simply reflect biophysical randomness, some degree of variability could carry functional significance. For example, it is possible that the immune system has evolved to leverage this heterogeneity to balance immune robustness with resource conservation.

### S2* cells’ quasi-digital response to immune stimulation

While the single-cell genomics studies^19–21^ are useful examples of *Drosophila* immune response heterogeneity, they offer only a single time-point snapshot, limiting interpretations to a binary, “all-or-nothing” view of cellular responses. In contrast, time-course studies in mammalian fibroblasts responding to TNF stimulation describe a “quasi-digital” mode of activation: initially, individual cells either fully activate NF-κB or not at all and increasing ligand dose both recruits more responsive cells and increases nuclear NF-κB amplitude and immediacy of the responders ^5,8^. At the other end of this spectrum, macrophages stimulated with LPS show widespread activation across the population, with some cells responding more strongly than others, suggesting an analog phenotype similar to that observed in snapshots of fly fat body cells ^8,21^.

Our results with S2* cells align more closely with the graded, quasi-digital model: a consistent subset of cells remains unresponsive, while increasing stimulus concentrations elevate the proportion of highly responsive cells. This may be a strategy to support varying functional outcomes in response to a range of stimuli. For example, the subset of cells displaying Immediate and Immediate with Decay behaviors may selectively activate a set of target genes that require a sudden high concentration of NF-κB to activate. However, the faster Relish attenuation rate of the latter limits transcription output, resulting in faster deactivation of signaling. This mode of response heterogeneity has important implications for infection *in vivo*, where pathogen concentrations can be highly variable throughout a tissue. Activation of only a subset of cells minimizes bystander damage in the inflammatory process, while a tunable NF-κB and transcriptional response in activated cells enables a stronger, localized immune response^8,9,18,22^.

### Nuances in Relish-driven transcriptional dynamics

As a starting point, transcription is frequently modeled as a two-state system, in which a promoter can be ON and producing transcripts or OFF and transcriptionally silent ^51^. If the transitions between these two states are controlled by single rate limiting reactions, the model predicts that the distribution of ON and OFF times will be exponential. We find non-exponential ON and OFF periods, suggesting additional model features are needed to recapitulate our observed transcriptional dynamics (Figure 6, S8A). A study of the dynamics of synthetic promoters in *Drosophila* embryos found that promoters with only TATA box motifs drove dynamics consistent with a two-state model, while promoters with an Inr motif or both an Inr motif and TATA box together drove dynamics consistent with a three-state model. In this case, the third state was consistent with promoter-proximal polymerase pausing prior to active elongation ^37^. Other work has suggested the addition of a refractory state after an ON period, in which a promoter is reset prior to subsequent activation, yielding non-exponential waiting times. The refractory state encodes temporal “memory” of prior activation, and a three-state model with a refractory period recapitulates NF-kB transcription in mammals ^52^. Our measurements do not discriminate between these possible elaborations to the two-state model. The promoters of *Dpt* and *Mtk* do contain both Inr and TATA motifs, but existing experimental data ^58^ did not suggest the promoters were paused, and a molecular source of a refractory state in this setting is also unknown. Characterizations of RNA Polymerase II pausing and enhancer output under different experimental conditions may reveal the mechanisms that underlie our observations.

We also note, as in previous work, that the portion of transcriptionally active cells decreases more quickly than the Relish nuclear fraction ^52^ (Figure 4A). Although we can predict the presence of transcription bursts using the Relish nuclear fraction (Figure 5C), we suspect one source of false predictions of a transcription burst is the decreasing probability of transcriptional activation over time. There are several possible mechanisms that could explain this apparent decrease in Relish’s ability to induce transcription. First, there may be an increase in the concentration of negative regulators of Relish-driven transcription as immune stimulation persists. A number of negative regulators of Relish transcription are known, including Charon (formerly known as Pickle), AP-1, Stat92E, or DSP1 ^56,57,59^. Charon inhibits the active, processed form of nuclear Relish from homodimerizing and binding to κB domains, potentially through the recruitment of the histone deacetylase dHDAC. AP-1, Stat92E, and DSP1 work together to repress Relish-driven transcription over the course of sustained immune stimulus. Second, dephosphorylation of Relish’s serines 528 and 529 would not affect Relish’s nuclear localization, but would impair its ability to drive transcription ^16^. Lastly, there may be other changes in cell state or cycle that suppress transcriptional activity more generally. Future studies may be able to distinguish these hypotheses with targeted gene knockdowns and repeated stimulation of the cells.

Our findings begin to bridge a critical gap in our understanding of the evolutionary strategies underlying host defense in the ancestral innate immune system. We demonstrate that, even in the absence of adaptive immunity, *Drosophila* cells demonstrate signaling heterogeneity and enhancer-driven tuning, which may allow them to mount graded innate responses to immune challenges. This suggests a conserved strategy for balancing rapid defense against metabolic cost and inflammation risk. Leveraging single-cell measurements across multiple pathway nodes, our work is the first to demonstrate the efficacy of both the HaloTag and RhoBAST systems to respectively track protein and transcriptional dynamics in live *Drosophila* cells, which has broad potential for future applications. These findings enable future work to build quantitative, predictive models of tissue-wide immune responsivity, incorporating gene promoter and enhancer architecture, upstream NF-κB dynamics, and chromatin environment.

## Methods

### Cell culture

Drosophila Schneider (S2*) cells obtained from the Steve Wasserman lab were grown in complete media containing Schneider’s Drosophila Medium (Gibco) supplemented with 10% Fetal Bovine Serum (Corning), 2mM L-glutamine (Gibco), 1% Amphotericin B (Gibco), and 50 U/mL Penicillin-Streptomycin (Gibco) at 28°C. Cells were passaged every 3-5 days.

### Plasmid generation

The attB sequence-flanked Relish reporter cassette *(gypsy -gypsy-pCopia-Blast-pIEX-Halotag-Relish)* was constructed via Gibson assembly of the gypsy insulator coding sequence amplified from S2* cell cDNA, *pAct5-eGFP-Relish* construct (gift from the Edan Foley lab), and 891 bp HaloTag sequence (gift from the John Ngo lab).

The dual Relish and transcriptional reporter was constructed by cloning AMP enhancer sequences (*Dpt*, *Mtk*, or *BomS2*) upstream of the *Drosophila* Kozak consensus sequence ^60^, lacZ coding sequence, and 8xbiRhoBAST RNA aptamer ^24^ (1248 bp sequence provided by the John Ngo lab). This transcriptional reporter was inserted between the *gypsy* insulator sequences of the Relish reporter to sequester enhancer activity ^36^. Complete plasmid sequences are available upon request.

### Cell line generation

A landing pad was integrated into the 25C6 locus on the second chromosome of the *Drosophila* genome as described previously ^29^. Ribonucleoprotein (RNP) complexes were assembled by combining purified Cas9 protein (IDT) with crRNA:tracrRNA guide RNA duplexes targeting pre-optimized cut sites at the target locus, using the previously-published guide RNA sequences^29^. RNP complexes were electroporated (MaxCyte) into S2* cells along with a repair cassette (*pIEX-Puro-T2A-eGFP)* flanked by attP sites and homology arms corresponding to the cut sites. Stably integrated cells were selected in complete media supplemented with 10 µg/mL Puromycin (Invivogen) for 10 days.

Following successful attP integration confirmation via eGFP expression and PCR verification, recombinase-mediated cassette exchange was performed by electroporating cells with attB-flanked reporter cassettes and a plasmid expressing PhiC31 integrase (Drosophila Genomics Resource Center #1368). RMCE positive cells were then selected using complete media supplemented with 25 µg/mL blasticidin (Corning).

### IMD stimulation and live-cell confocal imaging

S2* cells were seeded in tissue culture 6-well plates (Corning) at 10^6^ cells/mL in complete media with blasticidin and incubated at 28 °C for 18 hours. Cells were then treated with 1 μm 20-hydroxyecdysone (Sigma-Aldrich) and incubated for 24 hours.

The following day, glass-bottom 8-well µ-Slides (iBidi) were rocked with 0.5 mg/mL concanavalin A (Sigma-Aldrich) for 30 minutes, washed 3× with water, and dried. 2.5 × 10^6^ cells/mL were seeded in the wells with complete media with blasticidin, 10 µg/mL Hoechst 33258 (Thermo) and 200 nM JFx650 (Tocris) and left to adhere for 1 hour. Wells were then gently washed 3× for 5 minutes with media and replaced with complete media with blasticidin, 2 µg/mL Hoechst 33258 (Thermo Scientific), 40 nM JFx650 (Promega), and 1× SpyRHO555 (Cytoskeleton) (imaging media).

After a set interval of baseline signal acquisition, imaging was paused and wells were injected with either 0, 1, 10, or 100 µg/mL of purified peptidoglycan isolated from *Bacillus subtilis* (Sigma) in imaging media at 20% of the well volume, and then imaging continued.

Cells were imaged with a Nikon Ax point scanning confocal microscope with high sensitivity GaAsP PMTs and 60× oil immersion objective (NA = 1.4), controlled by Nikon Elements software. The same laser and acquisition settings were used for each set of experiments (see below). All the experiments were repeated at least thrice to ensure reproducibility. For imaging the Relish reporter without enhancers, frames were captured every 10 minutes for 1.5 hours, then every 30 minutes for 13 hours. Sparse imaging of the Relish and transcriptional reporter had acquisition every 15 minutes for 2 hours, every 30 minutes for 8 hours, then every hour for 5 hours. Dense imaging was done with imaging every 30 seconds in two hour intervals. Detection of RhoBAST aptamer transcription is delayed approximately 2 minutes, due to the time required to transcribe the 3058 bp 3’ lacZ sequence ^41^.

All images were acquired with a 2 µm z-stack over 3 frames. Hoechst 33258 stained nuclei were imaged with a 405 nm laser at 0.3% intensity and 50 gain, with an acquisition range of 429-474 nm. JFx650 labeled Relish was imaged with a 640 nm laser at 0.1% intensity and 30 gain, with an acquisition range of 652-712 nm. SpyRHO555 labeled RNA was imaged with a 561 nm laser at 1% laser power and 40 gain, with an acquisition range of 571-625 nm. Images were acquired as 1024×1024 Nikon ND2 files, and experimental replicates were imaged on different days.

### Data pre-processing and segmentation

Cell image slices were maximally projected along the z-axis using Fiji^61^ and time courses converted to TIF files. Cell and nuclei segmentation and tracking were performed using a custom script automating Fiji’s Trackmate plugin ^61,62^. Segmentation was done with Cellpose v2.2.3 using a custom-trained cell body model and pretrained nuclear model ^62,63^. Cell diameter was set to 10 um and the default value of 0.4 for flow threshold led to robust segmentation. Cell and nuclear masks with areas less than 40 µm^2^ and 8 µm^2^, respectively, were removed. The LAP tracker was used for tracking of segmented cells over time ^64^ with a max gap frame = 15, max gap closing distance = 10 pixels, and linking max distance = 5 pixels. Track splitting and merging were not allowed.

Cell and nuclei masks were interpolated over time to fill in missing frames with a custom Python script. Nuclei masks were then subtracted from cell masks to form cytoplasm masks, corresponding cytoplasm and nuclei label values were matched, and cells without nuclei were removed from the dataset ^65^.

For segmentation of SpyRHO555-labeled RhoBAST transcriptional foci, we implemented an ilastik-based workflow combining interactive machine learning with automated pixel classification^66^. Using ilastik’s Pixel Classification workflow, sparse annotations were manually drawn labeling nuclei, RhoBAST foci, and background regions on representative frames. A Random Forest classifier then iteratively learned decision boundaries in the feature space using these annotations, and regions with conflicting probabilities received additional annotations to refine class separation. The trained classifier generated per-pixel probability maps (nuclei, foci, background), which were then processed through ilastik’s Object Classification workflow to generate binary foci masks. Simple thresholding of 0.25 and a maximum size filter of 200 were applied to minimize false positives. This classification pipeline was then applied batch-wise to all timepoints across all transcriptional imaging files, and the resulting segmentation masks were exported as TIF stacks for downstream tracking analysis.

All cells underwent manual verification of their cytoplasmic and nuclear masks against the dye labels before being added to the dataset based on the following criteria:

Inclusion criteria (must hold in every frame):

1. Entire cell body is in focus and fully within the frame
2. Cell displays healthy morphology (intact body and consistent movement)
3. Cytoplasmic and nuclear masks precisely match their dye-labeled outlines

Exclusion criteria (if observed at any point):

1. Cell moves out of view
2. Cell dies (sudden shrinkage or loss of movement)
3. Cell divides
4. Either nuclear or cytoplasmic mask deviates from its designated region

### Relish and transcriptional foci quantification

A custom Python script was initialized with multichannel TIFFs for each cell (nuclei, JFx650-labeled Halotag-Relish, and SpyRHO555-labeled RhoBAST (when applicable)), as well as masks for cell cytoplasms, nuclei, and transcriptional foci. Nuclear and cytoplasmic masks were applied over the Halotag-Relish channel, and R_nuc:tot_ was calculated for each cell over each time frame by dividing the nuclear Relish signal by the sum of the nuclear and cytoplasmic signal. Fold-change R_nuc:tot_ was calculated by normalizing the full R_nuc:tot_ time trace by the average value of pre-stimulus frames. Traces were smoothed over time with Savitzky Golay filtering with a polynomial order of 2.

For transcriptional data, foci were first filtered to only include those detected within nuclei, then the sum of the RhoBAST signal across the segmented pixels (Int_RB_) was calculated for each detected focus. No smoothing was performed for sparse imaging analysis. Dense imaging transcriptional data was smoothed over time with Savitzky Golay filtering with a polynomial order of 2. Python’s find_peaks function was applied to smoothed traces for detection of transcriptional bursts with a minimum prominence of 200 units, and burst durations were calculated at a relative height of 0.9. Burst start times were shifted 2 minutes earlier due to time necessary for lacZ transcription^41^. Each cell’s average burst slope, and each individual burst’s on and off times were calculated.

### Support Vector Machine generation and training

#### Cell trace behavior classification of post-stimulus traces

Descriptors of post-stimulus smoothed cell traces for the fold-change Relish nuclear localization were extracted from single-cell data (1592/1592 cells, Figure 2B) using custom Python scripts. Those descriptors (per trace) included:

1. Maximum value (Max): Maximum value of the Relish nuclear fraction as measured by fold-change relative to the pre-stimulus average.
2. Time of maximum value (*t*_max_): Time at which the Relish nuclear fraction is maximal.
3. Time of half-maximum value (*t*_half_): Time at which the Relish nuclear fraction first rises to half the maximum value.
4. Area (AUC): Area under the trace integrated using trapz() from the scipy.integrate library in Python.

These data were manually scaled using min-max normalization (Eq. 1)

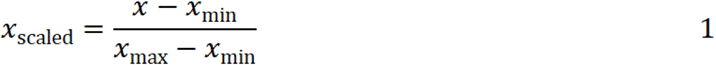

where *x* is the value of one of the four parameters above, with ranges *x*_min_, *x*_max_ of [1, 1.8] for Max; [0, 1000] for *t*_max_; [0, 500] for *t*_half_; and [900, 1500] for AUC.

We then manually classified a subset of traces (training dataset; 150/1592 cells) from the nuclear localization dataset (Figure 2B) into five distinct categories based on their short- and long-term response dynamics (Figure S3): Immediate (I), Immediate with Decay (Id), Gradual (G), Delayed (D), and Nonresponsive (N). We used a custom Python script to randomly retrieve individual traces for visual classification and selected the training dataset to include a representative sampling of all five behaviors.

We then used C-support classification (C-SVC) to build an SVM classification model (Classifier SVM A) taking as input the descriptor values for the cells in the training dataset. We used GridSearchCV() from the sklearn.model_selection library in Python to train and optimize our model via cross-validated search over a parameter grid. We used C-support vector classification with 5-fold cross-validation and scored for accuracy using accuracy_score() from the sklearn.metrics library in Python. The final optimized Classifier SVM A used a linear kernel with C, gamma, and degree values of 100, 0.1, and 2 respectively. We then used this model to classify the remaining cells (testing dataset; 1442/1592 cells) into the five behavior categories.

We retrained a second model (Classifier SVM B) to classify the traces from the transcriptional reporter dataset (3292 cells) (Figure 4A, S7) into the same five behavior categories following the same procedure.

#### Long-term behavior prediction of pre-stimulus traces

Descriptors of pre-stimulus cellular characteristics were extracted from single-cell data (1592/ 1592 cells) using custom Python scripts. Those descriptors (per cell) include:

1. Absolute cytoplasmic Relish intensity;
2. Absolute nuclear Relish intensity;
3. Absolute total (cytoplasmic and nuclear) Relish intensity;
4. Ratio of nuclear to cytoplasmic Relish intensity;
5. Absolute cytoplasmic cellular area;
6. Absolute nuclear cellular area; and
7. Absolute total (cytoplasmic and nuclear) cellular area.

For each of these descriptors, the pre-stimulus average, standard deviation (SD), and area under the curve (AUC) were calculated to yield a total of 21 features per cell. The data were scaled using MinMaxScaler() from the sklearn.preprocessing library in Python.

We then built two SVM models to predict a cell’s long-term behavior given its pre-stimulus cellular characteristics. Prediction accuracy was determined by comparison to the post-stimulus SVM classification results described above. For the predictor models, we collapsed the five behavior categories from the classification models into two: (1) immediate response (I), comprised of the previous Immediate (I) and Immediate with Decay (Id) categories; and (2) non-immediate response (NI), comprised of the previous Gradual (G), Delayed (D) and Nonresponsive (N) categories.

The first (Predictor SVM A) was trained on a dataset including all 21 features and was built and optimized as described above. The final optimized Predictor SVM A used a radial basis function (RBF) kernel with C, gamma, and degree values of 100, 0.1, and 2 respectively.

The second model (Predictor SVM B) performed recursive feature elimination (RFE) with cross-validation to select features using RFECV() from the sklearn.feature_selection library in Python. 5-fold cross-validation was done using StratifiedKFold from the sklearn.model_selection library in Python. RFE analysis was repeated 100 times to identify features appearing in at least 80% of the trials. The final set of optimized features included the nuclear Relish intensity AUC, and the nuclear:cytoplasmic Relish ratio SD and AUC. The SVM parameters were then optimized as described above. The final optimized Predictor SVM B used a linear kernel with C, gamma, and degree values of 100, 0.1, and 2 respectively.

### Transcription factor binding and promoter analysis

We obtained binding site motifs of known immune-responsive transcription factors, including those for Relish (motif obtained from Fly Factor Survey^67^ [FFS] and i-cisTarget^68,69^ [ict]), all five GATA factors, including Pannier, Serpent, Grain, dGATAd, and dGATAe (FFS, ict)^46^, kay/Jra (activated in the pro-inflammatory Jak/Stat pathway, FFS)^70–72^, EcR/USP/Eip74EF (activated via the Ecdysone pathway, FFS, ict)^73^, gcm (involved in immune responses in hemocytes, ict)^74^, and slp2/forkhead (required for circulating hemocyte differentiation, FFS)^75^. We then used the MEME suite’s FIMO tool ^44^ to independently search for the presence of these motifs in the sequences of *Dpt*, *Mtk*, and *BomS2* enhancers, using a match p-value < 1E-3.

For promoter analysis, we input enhancer sequences into the Elements Navigation Tool^42^ and identified key promoter elements across all enhancers, such as TATA boxes and initiator sequences.

### Statistical analysis

All statistical analyses were performed using Python’s scipy.stats library. χ² tests with false discovery rate (FDR) corrections were performed to compare Relish behavior category distributions across stimulation concentrations (Figure 2C, S7), Kolmogorov–Smirnov tests were performed to compare kernel density estimate distributions of tFoci+ and tFoci-metrics (Figure 5), DeLong’s tests with Bonferroni corrections were performed to compare the areas under the ROC curves (Figure 5), 2-sided Mann–Whitney tests with Bonferroni corrections were performed to compare burst parameter distributions (Figure 6, S8B) and Kolmogorov-Smirnov maximum likelihood estimation tests were performed to evaluate how well burst OFF times conform to an exponential model (Figure S8A). P-values of <0.05, <0.01, denoted as * and ** respectively, were considered statistically significant. All statistical tests’ resulting p-values are included in Supplementary Table 2.

## Acknowledgements

This work was funded by NSF Award 2223888 (to ZW) and NIH 5T32GM130546 (to NN). We thank Francis Vu for help in constructing an earlier version of the enhancer reporter constructs. We also thank John Ngo and Alex Green for helpful advice on imaging strategies, Tom Gilmore for discussion of NF-κB, and Nitin Kulkarni and Alfred Sloan from MaxCyte for help with cell electroporations. S2* cells were a gift from Steven Wasserman.

