## Supplementary Figures and Tables for "Heterogeneous NF-κB activation and enhancer features shape transcription in *Drosophila* immunity"

**Supplementary Figures, Tables, Video Captions**

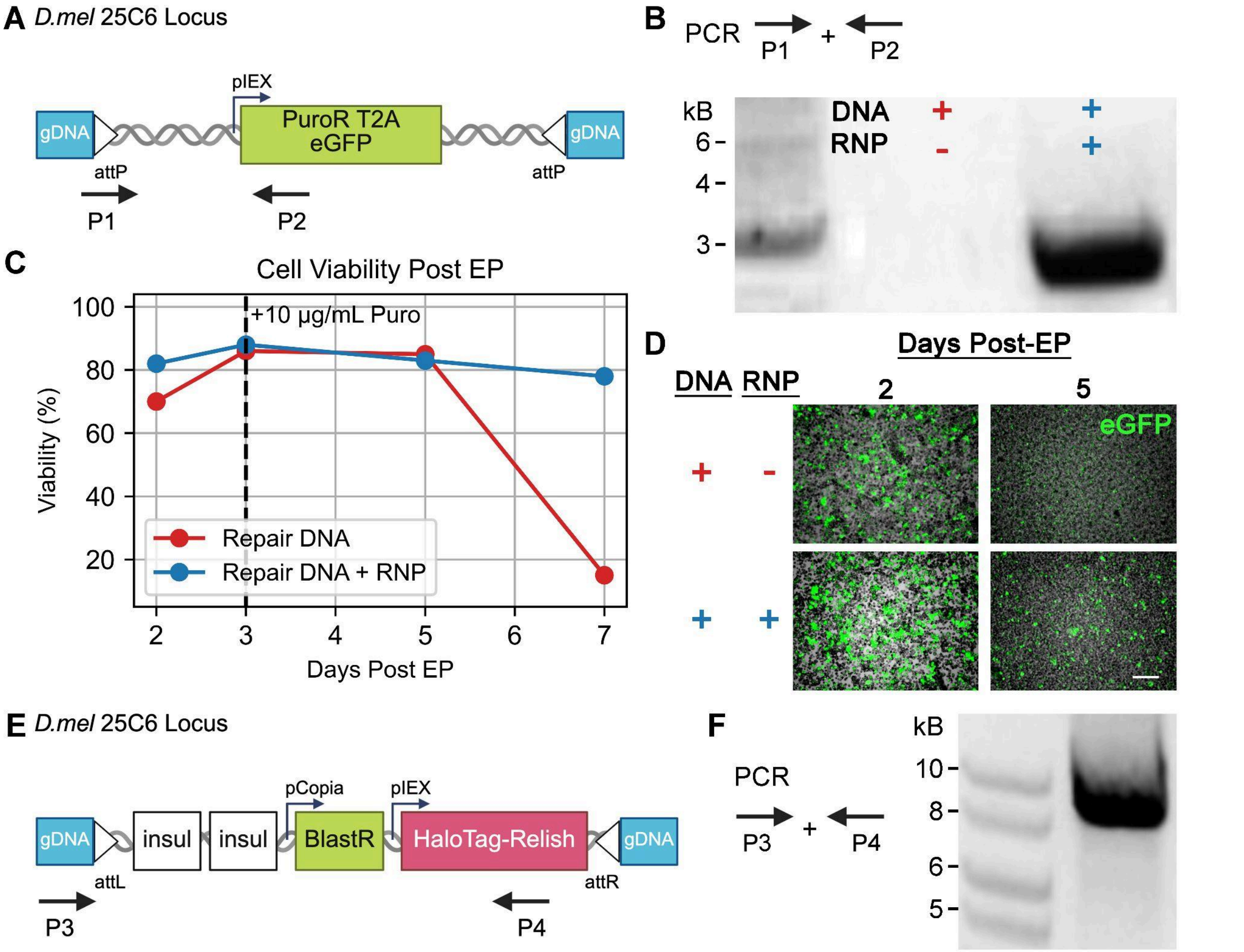


**Figure S1. Verification of CRISPR/Cas9-mediated integration of a landing pad at the 25C6 locus.**

**A.** Schematic of the DNA repair template inserted at the 25C6 locus with CRISPR Cas9 via homology-directed repair, containing puromycin resistance (PuroR) and eGFP flanked by attP sites. PCR primers P1 and P2 are indicated for genotyping (Supplementary Table 1). **B.** PCR genotyping of cell populations following electroporation (EP) with repair DNA alone (red, RNP–) or with both repair DNA and CRISPR/Cas9 ribonucleoprotein (blue, RNP+). A a product of expected size is observed only in the presence of RNP, confirming targeted integration. **C.** Quantification of cell viability over time following EP and puromycin selection. Co-delivery of repair DNA with RNP (blue) maintains higher viability compared to repair DNA alone (red). **D.** Representative fluorescence images showing eGFP expression at days 2 and 5 post-EP. 5 days post-EP, eGFP is only expressed in subpopulations in condition including RNP, indicating successful integration and expression from the landing pad. Scale bar = 100 μm. **E.** Schematic of the engineered 25C6 locus after site-specific ΦC31 recombinase-mediated cassette exchange to introduce gypsy insulators (insul), blasticidin resistance (BlastR), and HaloTag-Relish flanked by attL/R sites. PCR primers P3 and P4 are indicated for genotyping (Supplementary Table 1). **F.** PCR verification of the final engineered locus using P3 and P4 show a product of expected size only in the presence of RNP, confirming targeted integration. Together, these results confirm efficient CRISPR/Cas9-mediated insertion of a landing pad at the 25C6 locus and subsequent site-specific integration of the Halotag-Relish reporter construct.

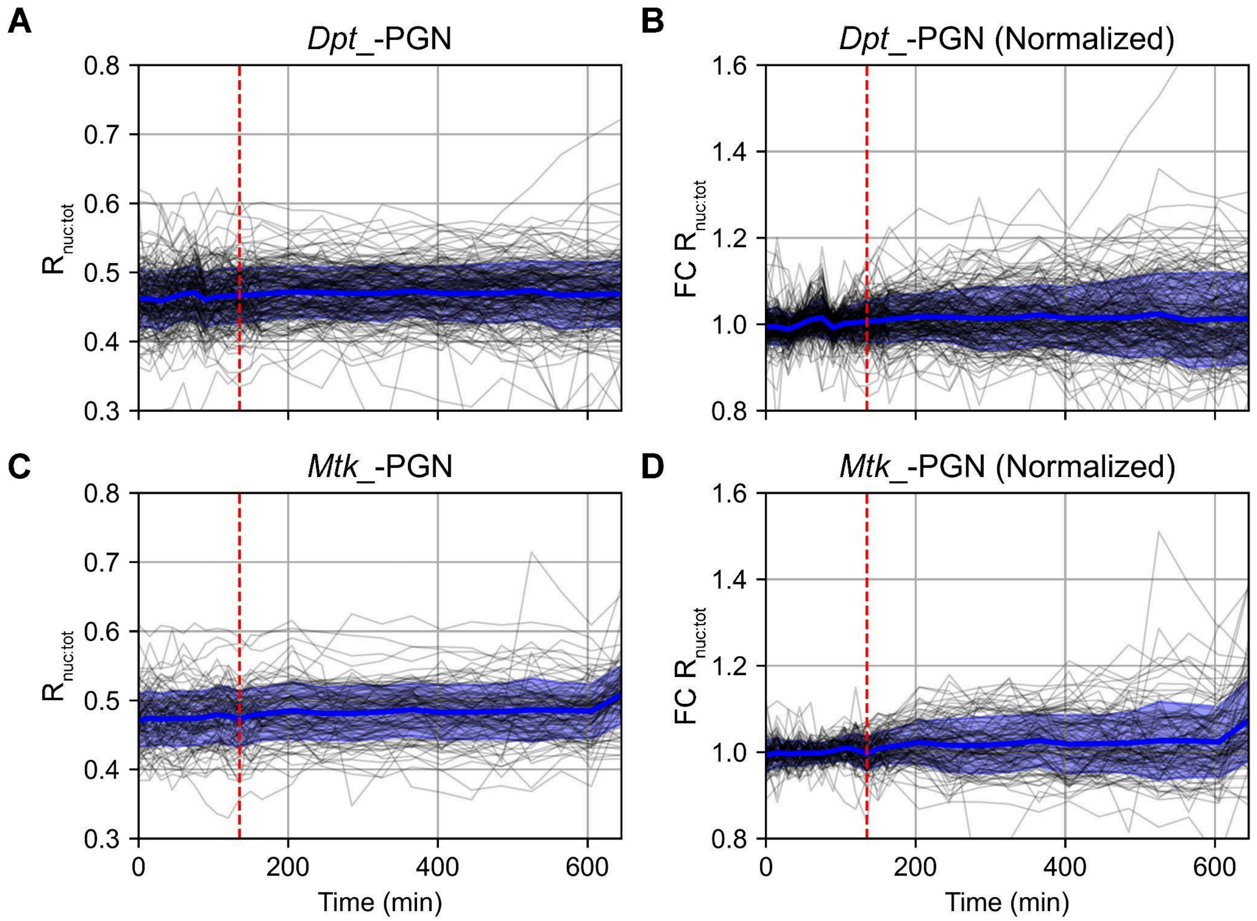


**Figure S2. Negative-control time courses of HaloTag–Relish dynamics in enhancer reporter lines upon water injection.**

**A.** Individual lines represent single cell $R_{nuc:tot}$ time courses and overlaid blue line and ribbon represent mean ± SD for the *Dpt*-enhancer reporter (n = 296 cells from 2 independent replicates). **B.** Corresponding fold-change in $R_{nuc:tot}$ after normalization to the mean pre-injection baseline (first 120 min) for the *Dpt*-enhancer reporter. **C–D.** Panels C and D replicate A and B, respectively, for the *Mtk*-enhancer reporter cells (n = 111 cells from 2 independent replicates) . In all panels, the vertical red dashed line denotes the time of water injection. A fluctuation of ±20% in both raw and normalized signals define the noise floor used to distinguish responsive from nonresponsive cells in Predictor SVM. Cells were imaged every 15 minutes for 2.5 hours, then every 40 minutes for 8 hours.

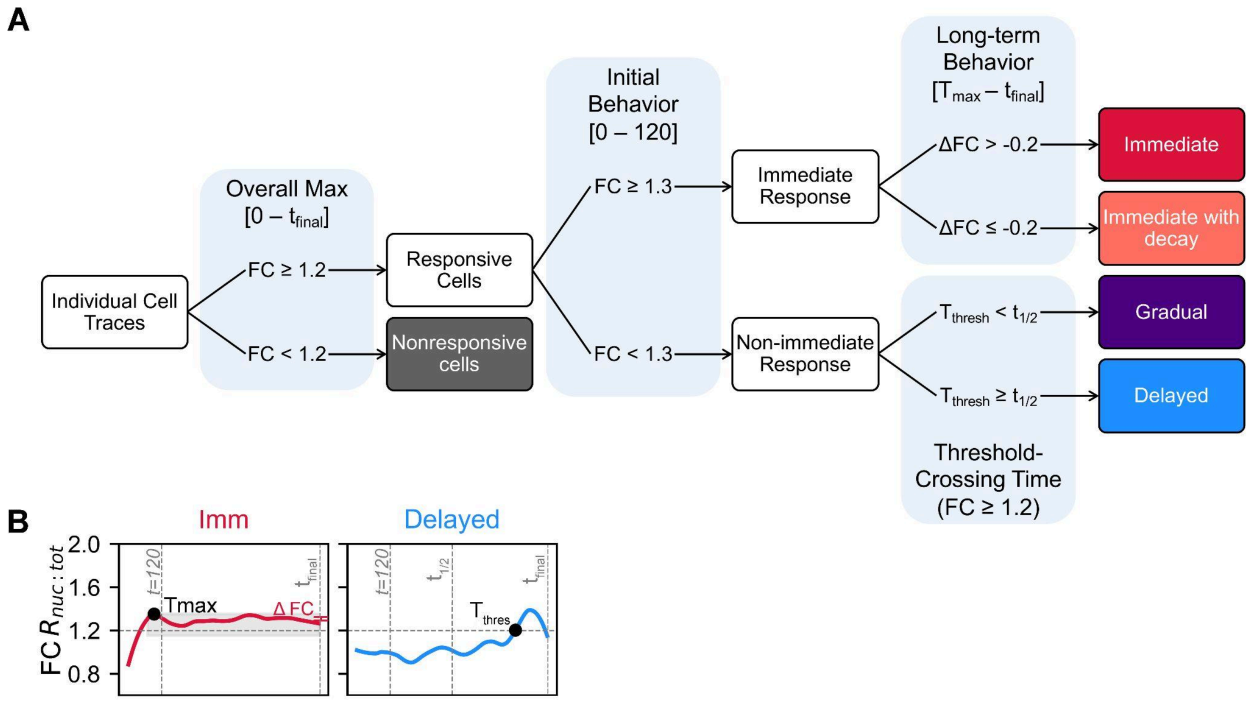


**Figure S3. Decision-tree workflow for assignment of single-cell Relish response classes for SVM training.**

**A.** Flowchart outlining heuristic rules applied to FC $R_{nuc:tot}$ traces to manually annotate dataset for Predictor SVM training. Categories included:

1. **Nonresponsive (N)**: never exceeds 20% increase (FC < 1.2) over the entire imaging period.

2. Responsive: $FC_{max}$ ≥ 1.2; then if FC ≥ 1.3 within 0–120 min → Immediate, otherwise → Non-immediate.

3. Immediate (I): peak FC ≥30% in the first 120 min. Subdivided by long-term decay: if ΔFC ($T_{max}$→$t_{final}$) ≤ –0.2 → **Immediate with Decay (Id)**, else remain **Immediate (I)**.

4. Non-immediate: time to first reach FC ≥ 1.2 ($T_{thresh}$) before $t_{1/2}$ (400 min) → **Gradual (G)**; within 400–800 min → **Delayed (D)**.

**B.** Example FC $R_{nuc:tot}$ traces for an Immediate cell (red trace; peak at $T_{max}$ <120 min, ΔFC measured to $t_{final}$) and a Delayed cell (blue trace; threshold crossing at $T_{thresh}$ >400 min). Vertical dashed lines demarcate the 120 min and 400/800 min decision windows; filled circles mark $T_{max}$ or $T_{thresh}$.

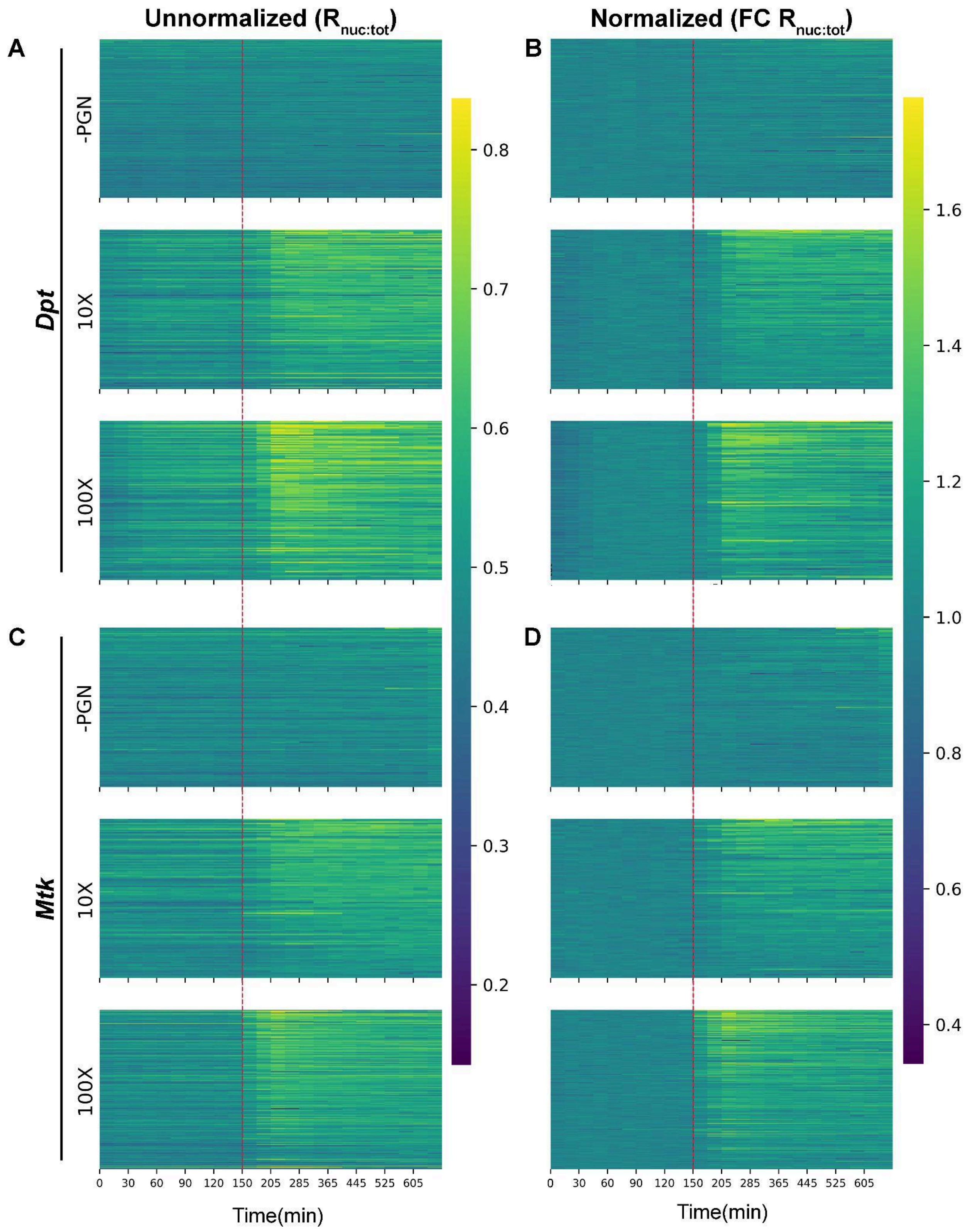


**Figure S4. Extended baseline imaging heatmaps of single-cell HaloTag–Relish intensity traces depicting pre-stimulus variability.**

**A.** Heatmap of raw nuclear HaloTag–Relish fluorescence intensity for Dpt-enhancer cells during pre-stimulus baseline followed by water (-PGN) injection (top, n = 196 cells), 10 µg/mL PGN (middle, n = 107 cells), and 100 µg/mL PGN (bottom, n = 94 cells) at t = 150 min **B.** Heatmap of fold-change in intensity after normalization to the mean baseline (first 150 min) for the *Dpt*-enhancer reporter cells. **C.** As in (A) for *Mtk*-enhancer cells (-PGN: n = 111, 10 µg/mL: n = 93, 100 µg/mL: n = 146 cells). **D.** As in (B) for Mtk-enhancer cells. Cells were imaged every 15 minutes for 2.5 hours, then every 40 minutes for 8 hours over two experimental replicates.

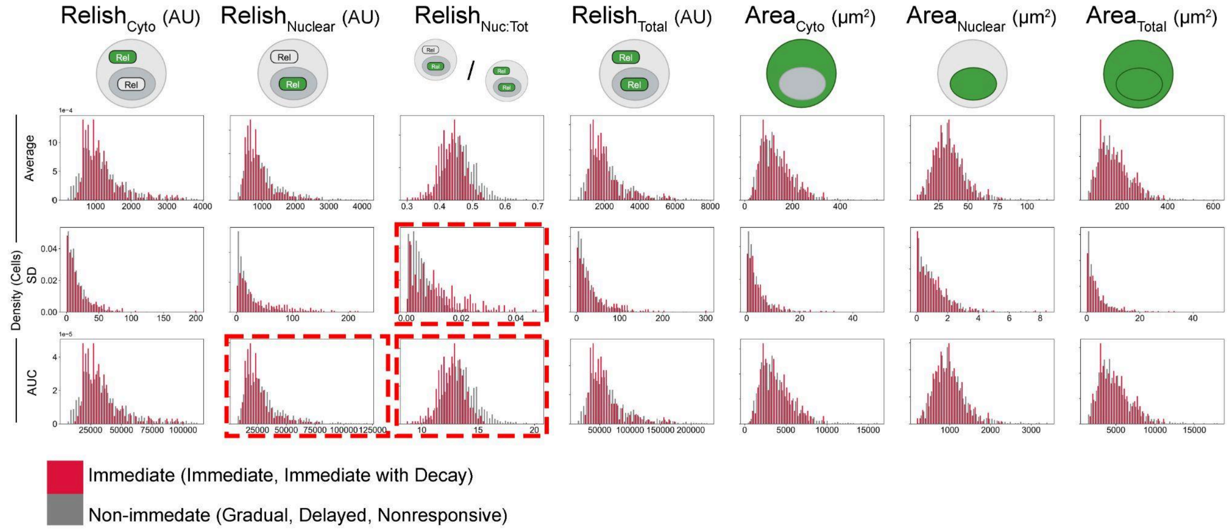


**Figure S5. Density histograms of pre-stimulus single-cell features used to train Predictor SVM.**

**A.** For each of seven pre-stimulus metrics measured (columns: cytoplasmic Relish intensity, nuclear Relish intensity, nuclear-to-total Relish ratio, total Relish intensity, cytoplasmic area, nuclear area, and total cell area), top row depicts schematic of measurement. Histograms of each metric's mean (second row), standard deviation (third row), and area under the curve (AUC, fourth row) over the 30 minute pre-stimulus baseline are shown (21 histograms total). Cells classified by the Predictor SVM as Immediate responders (red; Immediate and Immediate with Decay) are overlaid on non-immediate cells (gray; Gradual, Delayed, Nonresponsive), illustrating the pre-stimulus variability exploited for predictive classification. Training the Predictor SVM on all 21 pre-stimulus features achieved 93.7% classification accuracy. Using recursive feature elimination, we found that just three features (boxed in red) —nuclear-to-total Relish standard deviation, nuclear-to-total Relish AUC, and nuclear Relish AUC—were sufficient to maintain high performance, with the pared-down model yielding 89.4% accuracy.

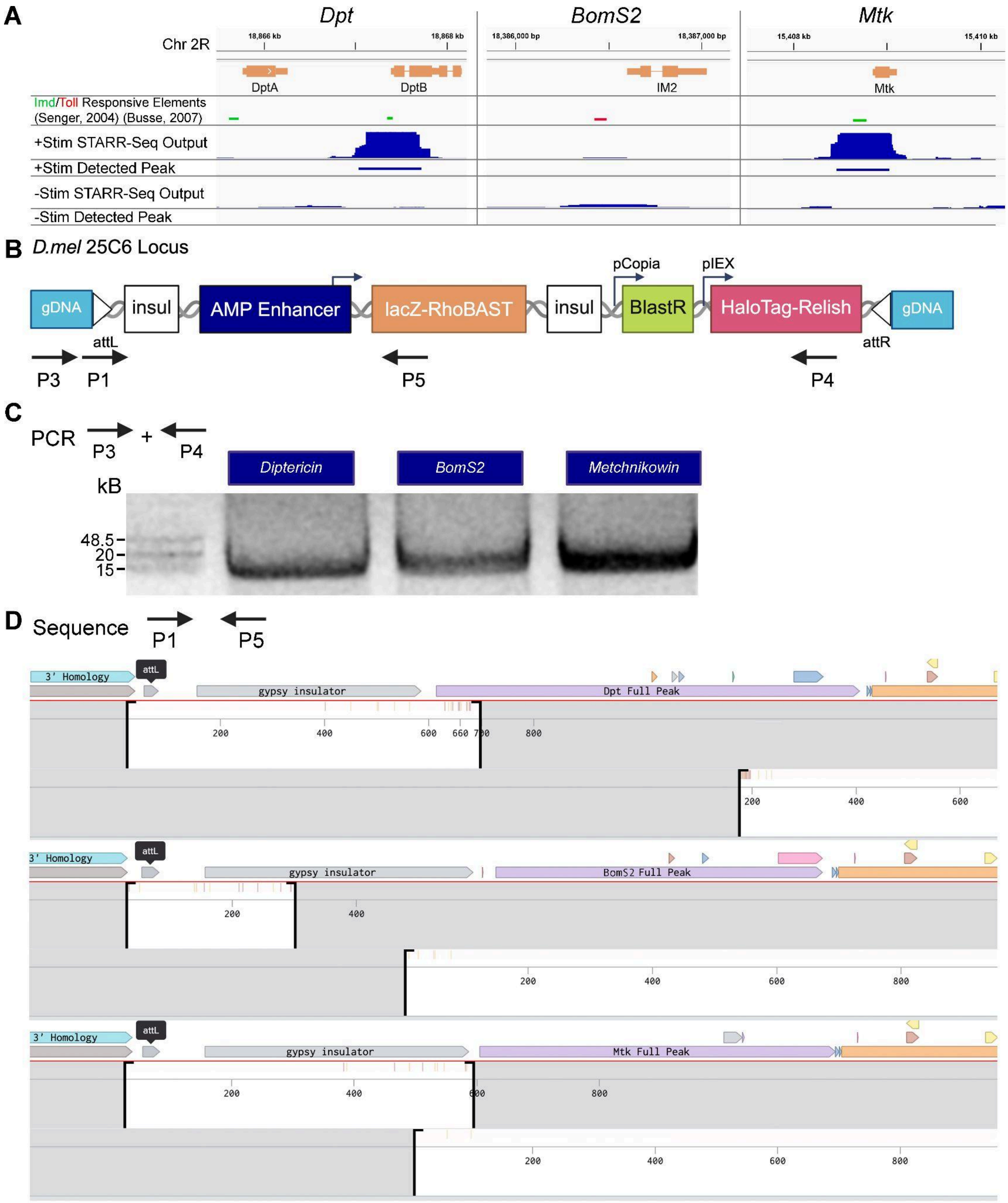


**Figure S6. Validation of *Dpt*, *BomS2*, and *Mtk* transcriptional reporter cell lines.**

**A.** For Dpt, BomS2 (also known as IM2), and Mtk AMPs (columns), the top row depicts published IMD/Toll (green/red bars, respectively) -responsive elements found at each loci. Below, STARR-Seq output signal and STARRPeaker peaks called based on output signal (indicated by the thick blue bars) at AMP loci in S2* cells with (+Stim, second and third row) or without (–Stim, fourth and fifth row) heat-killed *Serratia marcescens* presenting DAP-type PGN (manuscript in preparation). **B.** Schematic of the engineered 25C6 locus after RMCE to introduce gypsy insulators (insul), AMP enhancer (STARRPeaker peaks identified in A), lacZ‑RhoBAST, blasticidin resistance (BlastR), and HaloTag–Relish flanked by attL/R sites. Arrows P3/P4, and P1/P5 denote genotyping PCR and Sanger sequencing primer positions, respectively (Supplementary Table 1). **C.** PCR verification of each enhancer reporter cell line's engineered locus using P3 and P4 primers, showing a single band of the expected size (~15–20 kb) for each PCR reaction. **D.** Sanger‑sequencing of PCR product shows coverage across each inserted locus, confirming intact reporter assemblies. Coverage plots for *Dpt* (top), IM2 (*BomS2*, middle), and *Mtk* (bottom) show reads spanning the 3′ homology arm, attL site, gypsy insulator (P1), full STARR‑seq identified enhancer region, and partial lacZ sequence (P5). Sequence alignment done using Benchling (https://benchling.com).

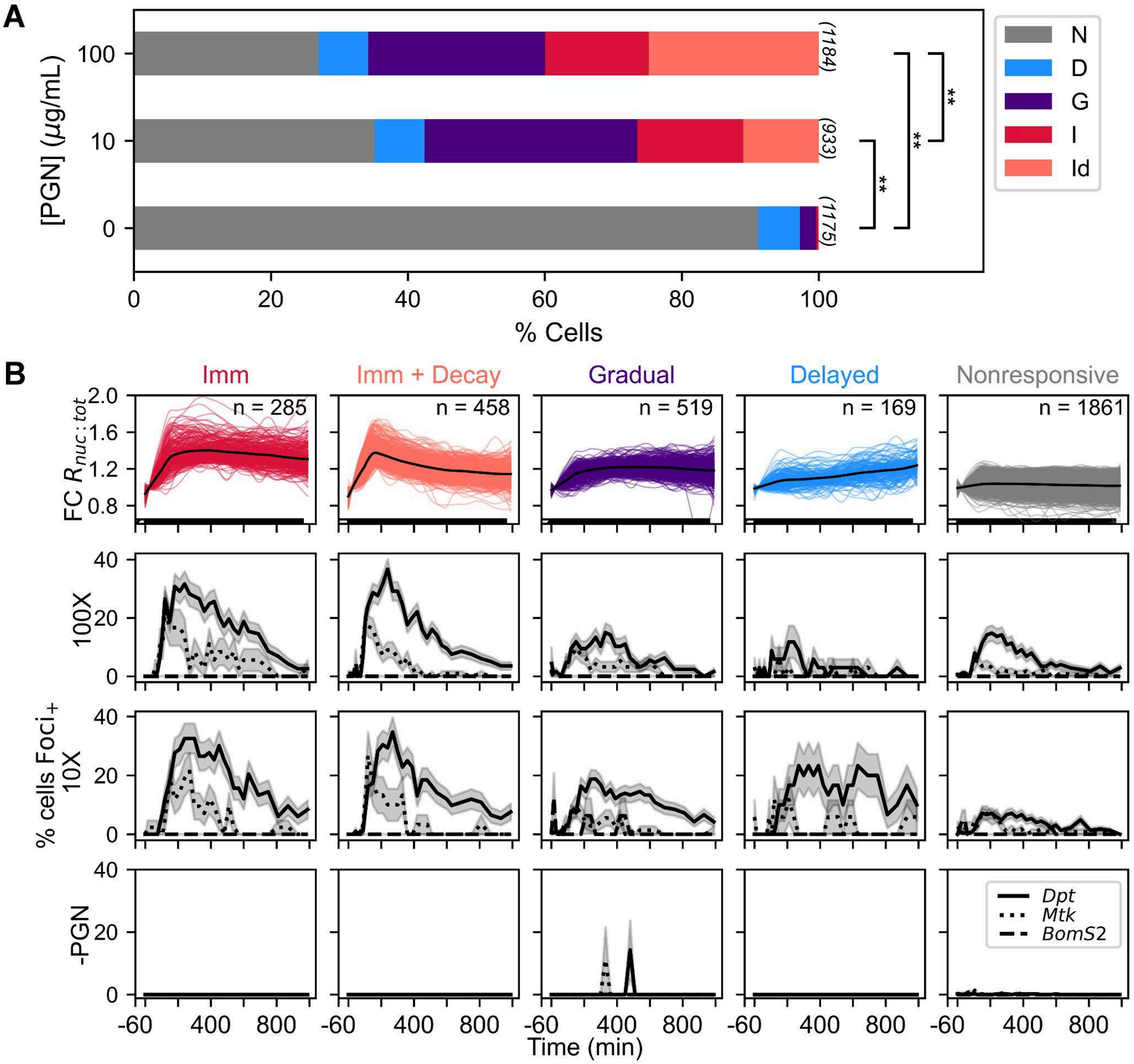

**Figure S7. Relish response behaviors and transcriptional foci dynamics across enhancer transcriptional reporter lines.**

**A.** Bar graphs depicting the percent of cells per Relish behavior category across PGN stimulation concentrations (n = 3,292 cells from 4 independent replicates). Cell counts for each condition are indicated. Comparable category distributions were obtained in HaloTag-Relish cells lacking a transcriptional reporter (Figure 2C). Significance was determined by a $\chi^2$ test with false discovery rate (FDR) correction. Corrected p-values for concentration pairs are included in Supplementary Table 2D (** $p < 0.01$). **B.** Transcriptional reporter cells stimulated with 100, 10, and 0 µg/mL PGN were categorized by the Classifier SVM (n = 3,292 cells, top row). Colored lines represent single cell traces and overlaid bold line represents the mean of each category. Of the cells within each category, the percent displaying transcriptional foci at each time point is plotted across reporters for *Dpt* (solid), *Mtk* (dotted), and *BomS2* (dashed) AMPs when stimulated with 100 (2nd row), 10 (3rd row), and 0 (4th row) µg/mL PGN. Traces represent mean ± SEM of all cells within category. Transcription dynamics follow similar trends across 100 and 10 µg/mL stimulus, but with generally more responding cells with the higher concentration. Minimal transcriptional activity is observed across all Relish behaviors for the *BomS2* reporter (dashed lines), and across all enhancers without PGN stimulus (bottom row).

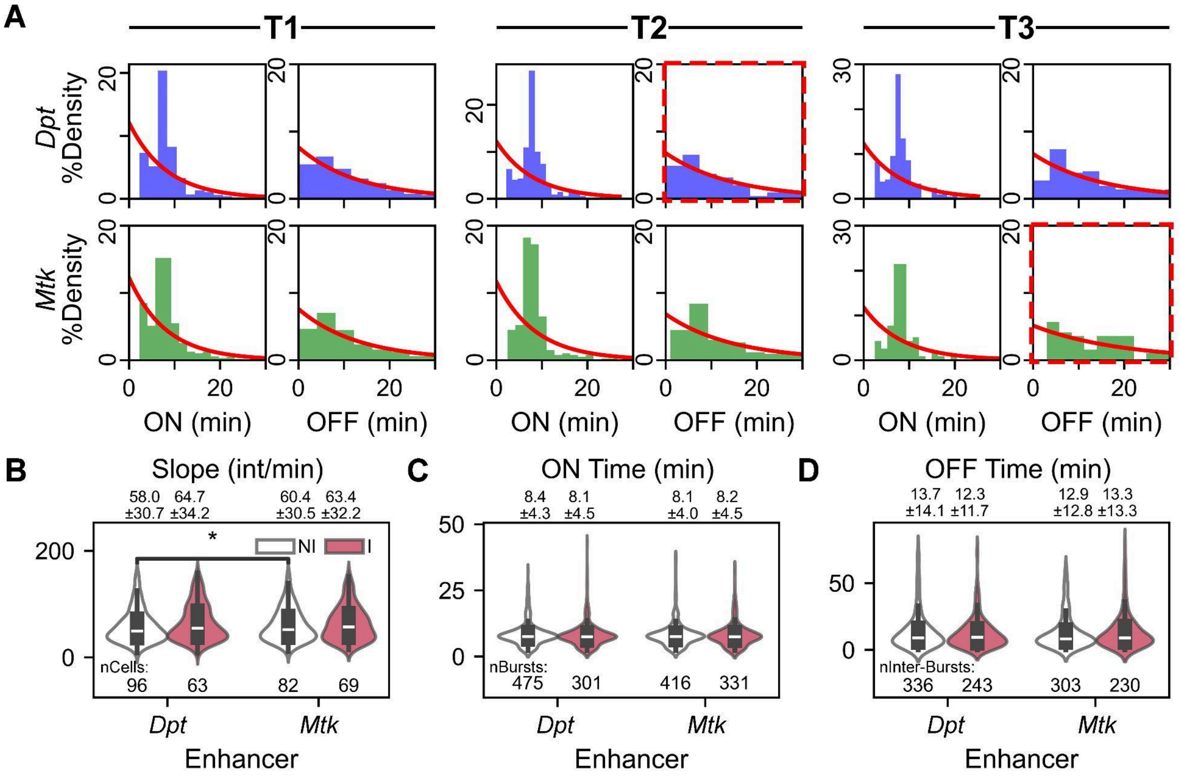


**Figure S8. Transcriptional burst ON/OFF duration distributions in *Dpt* and *Mtk* enhancer reporters.**

**A.** Probability density histograms of burst ON (left) and OFF (right) durations for *Dpt* (blue, top) and *Mtk* (green, bottom) reporters, measured in three post-stimulus windows each spanning 2 hours (Figure 3B). Red lines show exponential maximum-likelihood fits; only the distributions highlighted by dashed red boxes (*Dpt* OFF in T2 and *Mtk* OFF in T3) passed the Kolmogorov–Smirnov Maximum Likelihood Estimation (KS MLE) test ($p > 0.05$). Significant p-values included in Supplementary Table 2E. **B-D.** Comparison of burst parameters (as in Figure 6A) for cells in the T1 window classified as either Non-Immediate (NI, white) or Immediate (I, red) in Relish response: **(B)** Average slope of fluorescence increase for each bursting cell (top 90% of observed bursts plotted, intensity/min) (NI *Dpt* vs. NI *Mtk* p= 4.3e-02), **(C)** Burst ON durations, **(D)** Burst OFF durations for individual bursts. Violin plots (with median ± IQR shown as boxplots) with n values indicated below each category, and significance determined by a Mann–Whitney test with Bonferroni correction.

| | **5' to 3' Sequence** |
|---|---|
| Primer 1 | AGGGAACCGTTGACCTTGTAA |
| Primer 2 | GCCGTTTACGTCGCCG |
| Primer 3 | CGATTTTTCAAGTGGCGGTCGAGAGTG |
| Primer 4 | ATAAGGGCCCTTAGATTGGGTTAACCAGTAGGG |
| Primer 5 | CCGTAATGGGATAGGTCACG |

**Supplementary Table 1.** Primers used for reporter cassette assembly and Sanger sequencing.

**Supplementary Table 2.** Resulting p-values of various statistical analysis.
**A.** Figure 2: Results of $\chi^2$ comparisons of Relish behavior distributions across PGN concentration pairs in Relish reporter cells. P-values are adjusted with false discovery rate (FDR) correction for multiple comparisons.

| **[PGN] Pairs (µg/mL)** | **p-value** |
|---|---|
| 0 / 1 | 2.6e-13 |
| 0 / 10 | 2.0e-57 |
| 0 / 100 | 1.1e-72 |
| 1 / 10 | 8.6e-28 |
| 1 / 100 | 1.0e-44 |
| 10 / 100 | 1.2e-05 |

**B.** Figure 5**:** Results of Kolmogorov–Smirnov test comparing KDE distributions of metrics across tFoci+ and tFoci- frames for the *Dpt* and *Mtk* transcriptional reporters.

| **Metric** | ***Dpt* p-value** | ***Mtk* p-value** |
|---|---|---|
| $R_{nuc:tot}$ | 1.1e-194 | 3.5e-57 |
| FC $R_{nuc:tot}$ | 1.4e-151 | 1.2e-35 |
| Avg $R_{nuc:tot}$ [-30:t] | 1.2e-191 | 2.2e-44 |
| Avg $R_{nuc:tot}$ [-60:t] | 3.2e-185 | 9.5e-36 |
| Avg $R_{nuc:tot}$ [0:t] | 1.4e-20 | 0.02 |

Results of DeLong comparisons of ROC AUC values across metric pairs. P-values are adjusted with Bonferroni correction for multiple comparisons.

| **Metric Pairs** | ***Dpt* p-value** | ***Mtk* p-value** |
|---|---|---|
| $R_{nuc:tot}$ / Fold-Change $R_{nuc:tot}$ | 1.2e-05 | 4.1e-07 |
| $R_{nuc:tot}$ / Avg $R_{nuc:tot}$ [-30:t] | Insignificant | 2.9e-17 |
| $R_{nuc:tot}$ / Avg $R_{nuc:tot}$ [-60:t] | Insignificant | 2.6e-21 |
| $R_{nuc:tot}$ / Avg $R_{nuc:tot}$ [0:t] | 9.9e-114 | 9.2e-92 |
| Avg $R_{nuc:tot}$ [-30:t] / [-60:t] | 1.1e-03 | 2.5e-23 |
| Avg $R_{nuc:tot}$ [-30:t] / [0:t] | 5.8e-131 | 8.2e-92 |
| Avg $R_{nuc:tot}$ [-60:t] / [0:t] | 1.3e-143 | 2.2e-89 |
| $R_{nuc:tot}$ / Fold-Change $R_{nuc:tot}$ | 1.2e-05 | 4.1e-07 |

**C.** Figure 6: Results of two-sided Mann–Whitney comparisons of burst parameter values across imaging time windows. P-values are adjusted with Bonferroni correction for multiple comparisons.

| *Mtk* Burst Parameter | Time Window Pairs | p-value |
|---|---|---|
| Slope | T1 / T3 | 7.3e-03 |
| OFF Time | T1 / T3 | 9.2e-05 |
| | T2 / T3 | 5.0e-03 |

**D.** Figure S7. Results of $\chi^2$ comparisons of Relish behavior distributions across PGN concentration pairs in transcriptional reporter cells. P-values are adjusted with false discovery rate (FDR) correction for multiple comparisons.

| [PGN] Pairs (µg/mL) | p-Value |
|---|---|
| 0 / 10 | 6.0e-178 |
| 0 / 100 | 1.6e-236 |
| 10 / 100 | 2.8e-14 |

**E.** Figure S8. Results of Kolmogorov–Smirnov Maximum Likelihood Estimation test to fit burst OFF times to exponential model (significant when $p > 0.05$).

| Enhancer | Time Window | p-value |
|---|---|---|
| *Dpt* | T2 | 0.402 |
| *Mtk* | T3 | 0.065 |

**Supplementary Video 1. Sparse imaging of the *Dpt* enhancer reporter line revealed prominent foci of RhoBAST aptamer transcription within the nuclei of cells also displaying nuclear-localized Relish.**

Images were collected over 3 channels: Hoechst 33258 stained nuclei (left), JFx650 labeled Relish (middle), and SpyRHO555 labeled RhoBAST aptamers (right). Frames were acquired every 15 minutes for 2 hours (with 1 hour pre-stimulus baseline), every 30 minutes for 8 hours, then every hour for 5 hours. Images are confocal maximum-intensity projections (three z-steps spanning 2 µm).

**Supplementary Video 2. Dense imaging of the *Dpt* enhancer reporter line revealed complete RhoBAST bursts.**

Images were collected over 3 channels: Hoechst 33258 stained nuclei (left), JFx650 labeled Relish (middle), and SpyRHO555 labeled RhoBAST aptamers (right). Frames were acquired every 30 seconds in two hour intervals. Images are single confocal slices.
